# Structurally informed resting-state effective connectivity recapitulates cortical hierarchy

**DOI:** 10.1101/2024.04.03.587831

**Authors:** Matthew D. Greaves, Leonardo Novelli, Adeel Razi

## Abstract

Neuronal communication relies on the anatomy of the brain, yet it remains unclear whether, at the macroscale, structural (or anatomical) connectivity provides useful constraints for modeling effective connectivity. Here, we assess a hierarchical empirical Bayes model that builds on a well-established dynamic causal model by integrating structural connectivity into resting-state effective connectivity via priors. *In silico* analyses show that the model successfully recovers ground-truth effective connectivity and compares favorably with a popular alternative. Analyses of empirical data reveal that a positive, monotonic relationship between structural connectivity and the prior variance of group-level effective connectivity generalizes across sessions and samples. Finally, attesting to the model’s biological plausibility, we show that inter-network differences in the coupling between structural and effective connectivity recapitulate a well-known unimodal– transmodal hierarchy. These findings underscore the value of integrating structural and effective connectivity to enhance understanding of functional integration, with implications for health and disease.

**Significance statement:** To advance the understanding of how neuronal populations interact *in vivo*, it is essential to develop models that integrate neuroimaging modalities. Here, we show that integrating structural connectivity into dynamic causal modeling of resting-state effective connectivity substantially improves model evidence, yields reliable inferences, and demonstrates face and construct validity *in silico*. Furthermore, this integration reveals that structural connectivity’s influence on effective connectivity varies along an established unimodal–transmodal cortical hierarchy. This finding provides the first evidence of a network-dependent modulation of the relationship between structural and effective connectivity in humans. Against the backdrop of sustained and widespread interest in dynamic causal modeling, this study highlights the added value of integrating structural connectivity-based constraints, offering a more biologically grounded account of brain dynamics.

## Introduction

Computational modeling facilitates the mapping of both structural and effective connectivity in humans using *in vivo* magnetic resonance imaging (MRI) (1, 2). Structural (or anatomical) connectivity refers to a network of nerve tracts (bundles of axons), while effective connectivity refers to the directed, time-dependent influence that one neuronal population exerts on another (a construct inferred via the application of generative models to neuroimaging data). Although it is tempting to assume that neuronal populations primarily communicate via prominent nerve tracts detectable at the macroscale of MRI, understanding how structural connectivity constrains effective connectivity remains an open challenge (3).

Here, we examine a hierarchical empirical Bayes model in which—put simply—structural connectivity scales the variability (or noisiness) of resting-state (task-free) effective connectivity. At the first (subject) level, we utilize a well-established biophysical model—a dynamic causal model (DCM) (4, 5)—that describes how directed interactions between a network of unobserved neuronal populations cause changes in blood-oxygen-level-dependent (BOLD) signals obtained via functional MRI (fMRI). In this context, model inversion—inferring DCM parameters from observed data—involves generating synthetic data under different parameter settings and comparing model predictions to empirical observations, thereby enabling a move backwards from effects (observed data) to underlying causes (interregional neuronal communication, or effective connectivity) (6).

The hierarchical model utilized here assumes that to generate subject-level data, one first samples a vector of effective connectivity parameters from a distribution whose variance is shaped by group-level structural connectivity (right, Fig. 1A), adds subject-specific noise to these group-level parameters (middle, Fig. 1B), and finally uses the resulting subject-specific parameters in DCMs to generate synthetic data (right, Fig. 1C). In practice, we do not invert this full model jointly, but instead make use of standard Bayesian tools in a multi-step approach. First, each DCM is inverted separately, yielding a Gaussian posterior distribution over effective connectivity parameters for each subject. The subject-level posterior means furnish the data summaries for a second-level random effects (RFX) model (left, Fig. 1C), which treats them as random samples from a group-level distribution (first inset, Fig. 1B) whose variance is modulated by structural connectivity derived from tractography applied to diffusion-weighted MRI (dwMRI) data (Fig. 1A). After inverting this second-level model, the resulting group-level posterior serves as an empirical prior—estimated from the data—for re-evaluating subject-level effective connectivity analytically (second inset, Fig. 1B), enabling a refinement of first-level inferences under the second-level model (right, Fig. 1C) (7, 8).

**Fig. 1.**
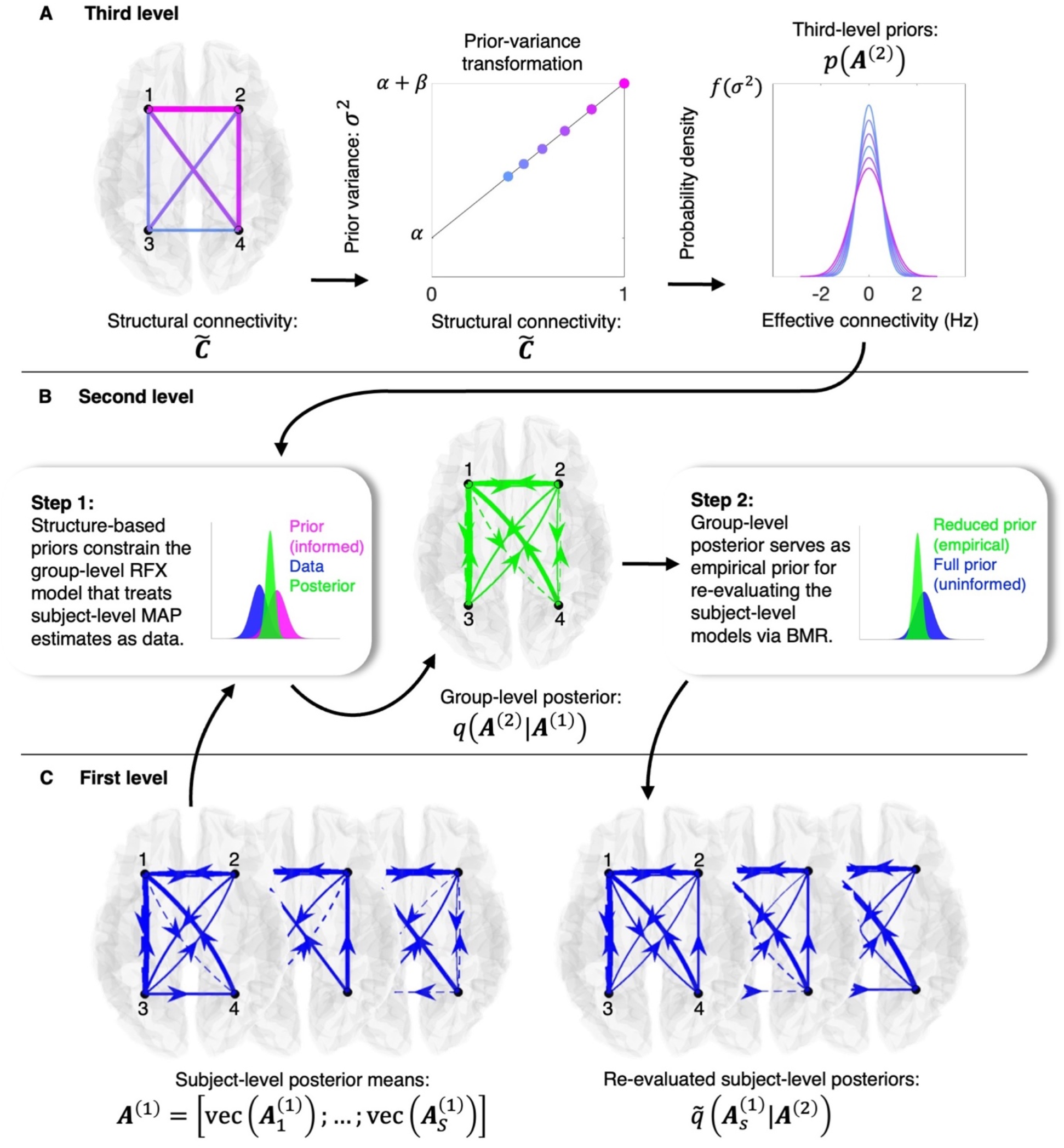
Inversion procedure for the three-level hierarchical empirical Bayes model. Arrows indicate the flow of information across levels. Inset text describes the procedure, and schematic, color-coded (univariate) densities indicate the level at which each (multidimensional) distribution originates: blue (first level), green (second level), and magenta (third level). Connectivity graphs use solid (dashed) lines to denote positive (negative) effective connections, with line weight proportional to connection strength. Notation is consistent with that used in the Methods. (*A*) At the third level, structural connectivity is transformed into connection-specific prior variances via a linear mapping 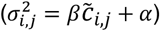 (Methods), which are used to construct priors over group-level effective connectivity parameters, *p*(***A***^(2)^). (*B*) At the second level, these structurally informed priors constrain the inversion of a random effects (RFX) model that treats subject-level posterior means—the maximum *a posteriori* (MAP) estimates—as data (Step 1). Note, however, that first-level posterior uncertainty is not discarded, but is retained through the Bayesian model reduction (BMR) within the hierarchical inversion. The resulting group-level posterior, *q*(***A***^(2)^|***A***^(1)^), then functions as a reduced (empirical) prior which, when combined with each subject’s full (uninformed) prior via BMR, enables analytic re-evaluation of the subject-level models (Step 2). (*C*) At the first level, each subject’s dynamic causal model (DCM) is inverted independently to yield subject-specific posterior means 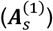, which are stacked and passed to the second level. Following second-level inversion, subject-level posteriors are re-evaluated as 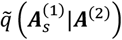.

Using *in silico* analyses, we show that this hierarchical model successfully recovers ground-truth effective connectivity and compares favorably with a popular structurally informed multivariate autoregressive (MVAR) model (9). In exploratory analyses of empirical data, we find—in line with theoretical expectations—a positive, monotonic relationship between structural connectivity and the variance of effective connectivity across 17 different resting-state networks associated with Schaefer and colleagues’ brain atlas (10). In tests of reliability and validity, we establish that network-specific relationships between structural and effective connectivity generalize out of session and out of sample. Finally, attesting to the model’s biological plausibility and relevance, we explore inter-network differences in the coupling between structural and effective connectivity and show that these differences align with an established cortical hierarchy as quantified by the principal gradient of functional connectivity (11).

## Results

Inferring effective connectivity from data using the hierarchical empirical Bayes model—introduced formally in the Methods—involves updating priors, where the *a posteriori* expectation and uncertainty of the model parameters is determined by combining priors with a likelihood function (that describes how well the model’s output fits the data). Formally, this process relies on Bayes’ rule:

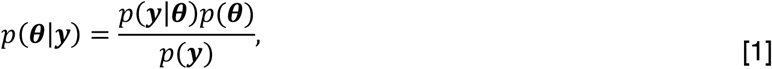

where *p*(***θ***|***y***) is the posterior probability of model parameters, ***θ***, under the observed data, ***y***, and the model. On the right-hand side, *p*(***y***|***θ***) is the likelihood of the data being observed under the model and its parameters, and *p*(***θ***) is the prior probability of the parameters under the model. These priors, *p*(***θ***), can be informative (for example, in this context, the prior variance of effective connectivity is assumed to scale with a linear transformation of normalized structural connectivity; Fig. 1A) or uninformative if little prior knowledge exists. So-called empirical priors are those which have been derived from the data itself (for example, when second-level models are used to re-evaluate first-level parameter estimates in a hierarchical framework; second inset, Fig. 1B).

Calculating the model evidence, *p*(***y***), which represents the marginal likelihood of the data under the model, requires integrating (or marginalizing) the likelihood over all possible values of the parameters, weighted by the prior distribution. In practice, computing this quantity is often infeasible, and thus, in this context, we make use of the well-known variational Bayes method under the Laplace approximation (VBL), which assumes that the true posterior distribution is Gaussian around its mode (6, 12). Assuming a Gaussian prior and likelihood, the fundamental VBL equation is:

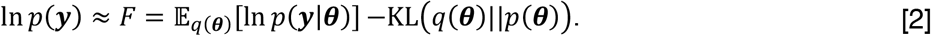

Here, *F*, is the (variational) free energy—a lower-bound on the log-model evidence—for the model, 𝔼_*q*(***θ***)_[ln *p*(***y***|***θ***)] is the expected log-likelihood under a variational—approximate posterior—distribution, *q*(***θ***), and KL(*q*(***θ***)||*p*(***θ***)) is the Kullback–Leibler divergence between the variational distribution, *q*(***θ***), and prior distribution. This formulation permits the optimization of *q*(***θ***) such that free energy is maximized, and the model optimally balances accuracy and complexity (6). For a broader introduction to these principles in the context of dynamic causal modeling, we refer the reader to Zeidman and colleagues (12).

Several of the results we present in this section reflect assessment of whether structural connectivity-based priors (henceforth referred to as structure-based priors) yield more parsimonious models (those that better balance accuracy and complexity). Conveniently, the free energy facilitates straightforward model comparison, where the log-Bayes factor is the difference between the log-model evidence assigned to a given set of observations under competing models. This permits comparison of alternative (inverted) hierarchical empirical Bayes models that differ, for example, in terms of whether a model, *m*_1_, incorporates structure-based priors, or whether the model, *m*_2_, incorporates uninformative priors:

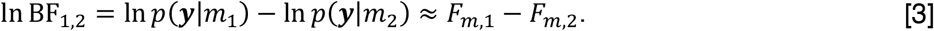

A positive log-Bayes factor, ln BF, indicates that the data provide more support for model *m*_1_ over *m*_2_, with larger values representing stronger evidence. A log-Bayes factor of 3 or higher is often considered strong— *e*^3^ ≈ 20 times more—evidence in favor of *m*_1_ (13).

For efficiency, we take an analytic approach to computing the approximate posterior (henceforth referred to as the posterior) and free energy for hierarchical empirical Bayes models with structure-based priors from an inverted model with uninformative priors (8). This analytic procedure, termed Bayesian model reduction (BMR) (14), is made possible due to the Gaussian form of Eq. 2. BMR enables one to adjust the sufficient statistics of the prior and obtain the free energy and posterior for a reduced (alternative) model from a full (parent) model analytically (without the need for optimization). Specifically, BMR yields the reduced model via the following relationships:

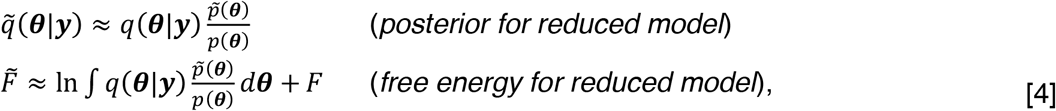

Where the tilde, (∼) indicates those elements relating to reduced models. Note that the likelihood terms do not appear in these expressions, as we assume the full and reduced models have identical likelihoods (allowing them to cancel out). For further details, we refer the reader to the Methods.

### Face and construct validity

#### In silico parameter recovery

In an *in silico* assessment of face validity, we generated 50 instantiations of BOLD-like time-series data for a 6-region effective connectivity network (SI Methods, Table S1–S2). In these simulations, group-level effective connectivity was derived from a standard Gaussian noise process scaled by a linear transformation of normalized structural connectivity, and subject-level effective connectivity was modeled as random deviations from group-level effective connectivity.

After inverting each DCM, the hierarchical model was inverted under uninformative priors (first and second insets, Fig. 1B). Then, focusing on the second-, group-level RFX model, we utilized a grid search and BMR to evaluate different parametrizations of the structural-connectivity-to-prior-variance transformation (henceforth referred to as the prior-variance transformation; middle, Fig. 1A) and score the resultant (structurally informed) reduced models against the (uninformed) full model in terms of the log-Bayes factor. We then applied Bayesian model averaging (BMA), weighting the hyperparameters governing each transformation by the resultant (normalized) model evidence (Procedures), thereby yielding an evidence-weighted prior-variance transformation. Finally, using BMR, the (uninformed) group-level model was re-evaluated under structure-based priors furnished via the evidence-weighted prior-variance transformation, and the posterior for the re-evaluated (structurally informed) group-level model was then treated as the prior under which the (uninformed) subject-level models were re-evaluated using BMR (right, Fig. 1C).

Fig. 2A and Fig. 2B show the normalized (sparse) structural connectivity network and group-level effective connectivity network utilized in this analysis, respectively. The parity plot (Fig. 2C) for the hierarchical empirical Bayes model indicates that—under a signal-to-noise ratio of 1—the model achieves good accuracy, with maximum *a posteriori* (MAP) estimates of group-level effective connectivity closely distributed around the identity line, a Pearson’s (product-moment) correlation (*r*) of 0.84, and root mean squared error (RMSE) of 0.181. Furthermore, Fig. 2D indicates that the evidence-weighted prior-variance transformation closely approximates the ground-truth variance of group-level effective connectivity.

**Fig. 2.**
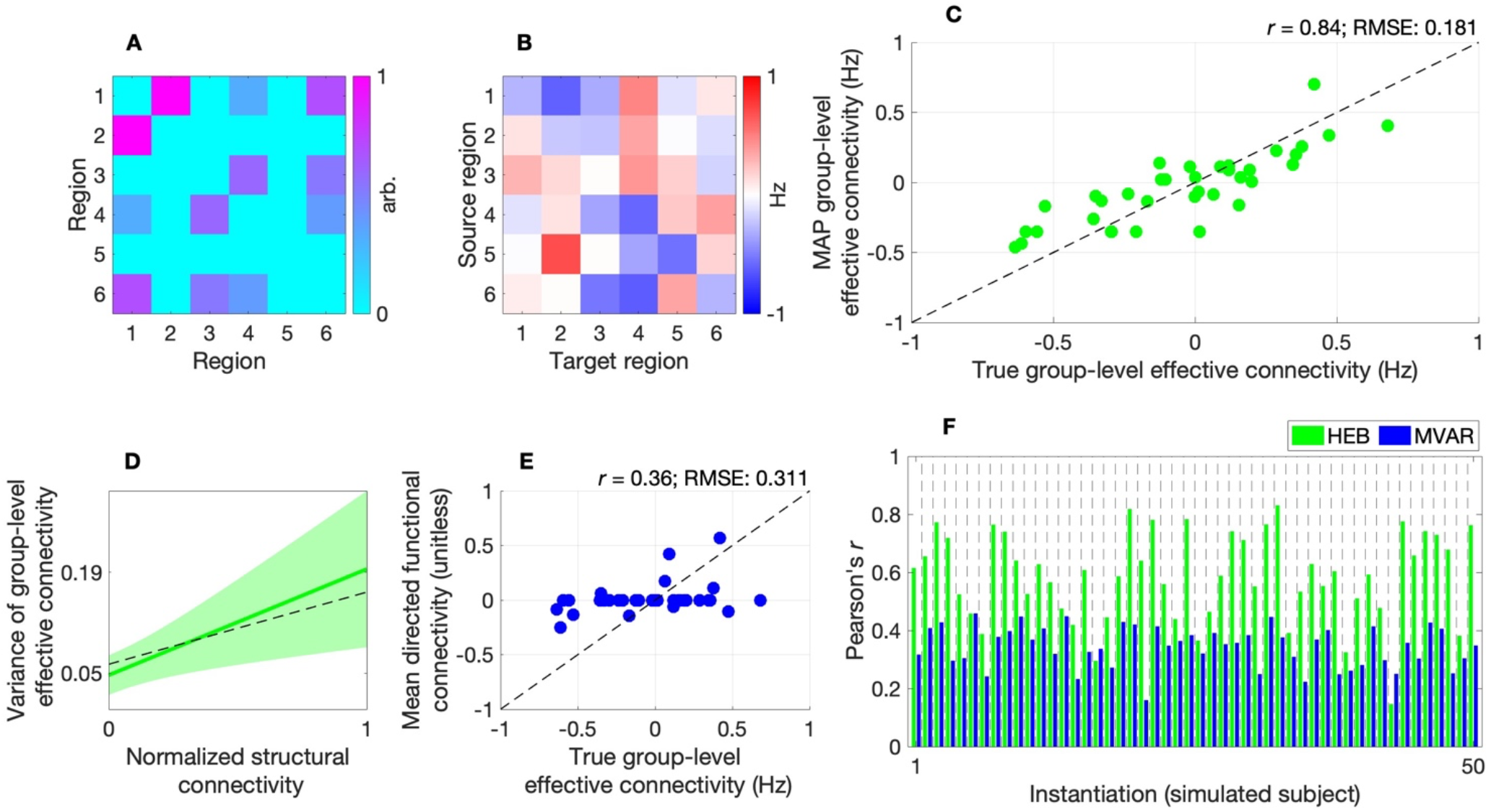
*In silico* evaluation of hierarchical empirical Bayes model. (*A*) Heatmap of normalized structural connectivity utilized in simulations. (*B*) Heatmap of group-level effective connectivity (derived from normalized structural connectivity) utilized in simulations. (*C*) Parity plot for the hierarchical empirical Bayes (HEB) model, showing maximum *a posteriori* (MAP) estimates of group-level effective connectivity plotted against true values. Scatter points are clustered around the (dashed) identity line with a Pearson’s (product-moment) correlation (*r*) of 0.84, and root mean squared error (RMSE) of 0.181, indicating good model performance. (*D*) Evidence-weighted prior-variance transformation (solid green line) with shaded area indicating 95% confidence envelope. The estimated transformation closely approximates the ground-truth variance of group-level effective connectivity (dashed black line). Y-axis tick marks indicate the transformation evaluated at the lower and upper bounds of the input parameter space. (*E*) Parity plot for the multivariate autoregressive (MVAR) model showing a lower correlation, higher RMSE, and increased zero-valued estimates. The mean directed functional connectivity is quantified via the mean of (unitless) autoregressive parameters across subject-level models. (*F*) Bar plot of correlation between true and estimated subject-level effective connectivity characterized via the HEB model (green bars) and MVAR model (blue bars) across 50 instantiations, highlighting the greater performance of the former model.

To provide construct validation, we show that the model outperforms a popular structurally informed multivariate autoregressive (MVAR) model of directed functional connectivity, for which structural connectivity serves as a template for possible interactions (such that directed functional connectivity is estimated from region *i* to region *j* if a direct structural connection exists between the two regions) (9). Relative to the hierarchical empirical Bayes model, this MVAR model showed lower accuracy at the group level (Fig. 2E) and tended to show poorer ability to recover effective connectivity at the subject level (Fig. 2F; SI Results, Fig. S1A–B). Finally, a macro F1-score demonstrates the superior ability of the hierarchical empirical Bayes model to classify positive, negative, and absent group-level effective connections across a range of noise conditions (SI Results, Fig. S1C).

#### Empirical parameter exploration

In an empirical face-validation phase we utilized both resting-state fMRI data—the session-1 recordings— and structural connectivity (obtained via tractography applied dwMRI data) from 100 healthy adults (54 female, age 22–35) sourced from the Human Connectome Project (HCP) (15). For each brain network in the Schaefer atlas, we inverted a separate hierarchical empirical Bayes model following procedures utilized in the preceding *in silico* analyses (Procedures). This enabled the derivation of an evidence-weighted prior-variance transformation per network.

Fig. 3A illustrates that, despite the evaluation of hyperparameter regimes resulting in flat prior-variance transformations (disregarding structural information), the resulting evidence-weighted prior-variance transformation for each network was a positive, monotonic function. Thus, models in which structural connectivity positively scaled with the prior variance of effective connectivity were, on average, the most parsimonious. The 95% confidence envelopes shown in Fig. 3A illustrate some uncertainty regarding the slope of these transformations and motivated the need to evaluate their reliability and out-of-sample validity.

**Fig. 3.**
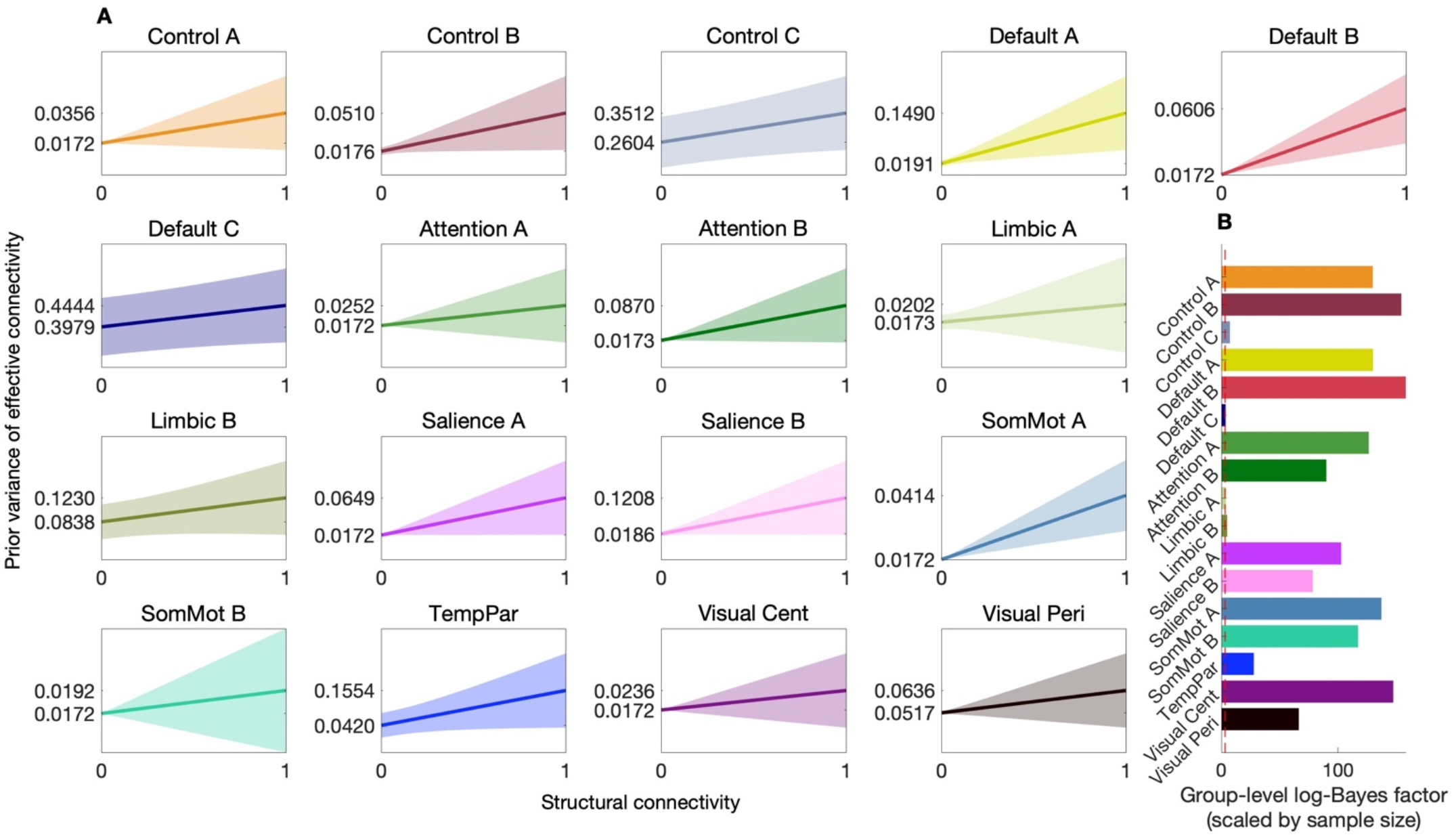
Bayesian model-averaged prior-variance transformation for 17 brain networks. (*A*) Each panel represents results for a hierarchical empirical Bayes model of a distinct effective connectivity brain network showing the evidence-weighted prior-variance transformation (solid line) as a function of normalized structural connectivity. The shaded regions indicate the 95% confidence envelope, reflecting uncertainty in the gradient of the transformations. Y-axis tick marks indicate the transformation evaluated at the lower and upper bounds of the input parameter space. Networks analyzed include control (Control A–C), default mode (Default A–C), attention (Attention A, B), limbic (Limbic A, B), salience (Salience A, B), somatomotor (SomMot A, B), temporoparietal (TempPar), and visual networks (Visual Cent, central; Visual Peri, peripheral). (*B*) Group-level log-Bayes factors (scaled by sample size) reflecting the per-subject increase in evidence seen when each hierarchical empirical Bayes model was inverted under its evidence-weighted prior-variance transformation. Note a dashed red line indicates log-Bayes factor of 3.

As depicted in Fig. 3B, across all networks, the increase in group-level evidence seen when each hierarchical empirical Bayes model was inverted under its evidence-weighted prior-variance transformation was substantial. At the group level, model evidence is obtained by summing subject-level free energies and applying a global complexity penalty (SI Methods). To better facilitate interpretation relative to standard Bayes-factor heuristics (13), we therefore report group-level log-Bayes factors scaled by sample size (henceforth referred to as sample-size-scaled log-Bayes factors), such that they reflect the average (or per-subject) evidence gain (14, 16). Under this metric, structure-based priors were consistently favored over uninformative priors across all networks, though with variability in effect sizes. In supporting analyses, we recovered a similar pattern of results—positive prior-variance transformations and increased evidence— using an alternative atlas (SI Results, Fig. S2).

### Test–retest reliability

Once network-specific evidence-weighted prior-variance transformations were identified using the session-1 dataset, we evaluated their test–retest reliability across time points—on the order of days—with hierarchical empirical Bayes models inverted using the session-2 data provided by the same subjects (Procedures). Specifically, we compared hierarchical empirical Bayes models with uninformative priors to hierarchical empirical Bayes models with structure-based priors furnished by applying the relevant, network-specific evidence-weighted prior-variance transformation (Fig. 3A). Results (SI Results, Fig. S3) indicate a consistent pattern across the networks, with substantially greater evidence for the models with out-of-session structure-based priors, and all MAP estimates of effective connectivity were in a plausible range.

### Out-of-sample validity

Next, we conducted two out-of-sample validations, assessing the degree to which the identified evidence-weighted prior-variance transformations served as a robust network-specific link between structural connectivity and prior variances. Utilizing both resting-state fMRI data—session-1 and -2 recordings—and structural connectivity from 50 healthy adults (24 female, age 22–35), out-of-sample validation mirrored the procedures utilized in the assessment of test–retest reliability. In this section, we present the out-of-sample validation results for session-1 data yet found similar results in the session-2 (SI Results, Fig. S4).

Fig. 4A indicates substantially greater evidence for the hierarchical empirical Bayes models that utilized the out-of-sample evidence-weighted prior-variance transformations, relative to models with uninformative priors. Note that for each network, we report the log-Bayes factor for both the group-level component of the model (semi-transparent bars), scaled by sample size, alongside log-Bayes factors for the subject-level component of the model (opaque bars), which are unaffected by this scaling. Under this metric, structure-based priors were consistently favored over uninformative priors across all networks. Control analyses in which prior covariance structures were randomly permuted suggest that such evidence gains are unlikely to result from the generic imposition of increased prior precision (SI Results, Fig. S5). We note too, that the introduction of these out-of-sample prior-variance transformations furnished MAP effective connectivity estimates within a plausible range (Fig. 4B), and that these effective connectivity estimates were highly consistent with those obtained in the other datasets (SI Results, Fig. S6).

**Fig. 4.**
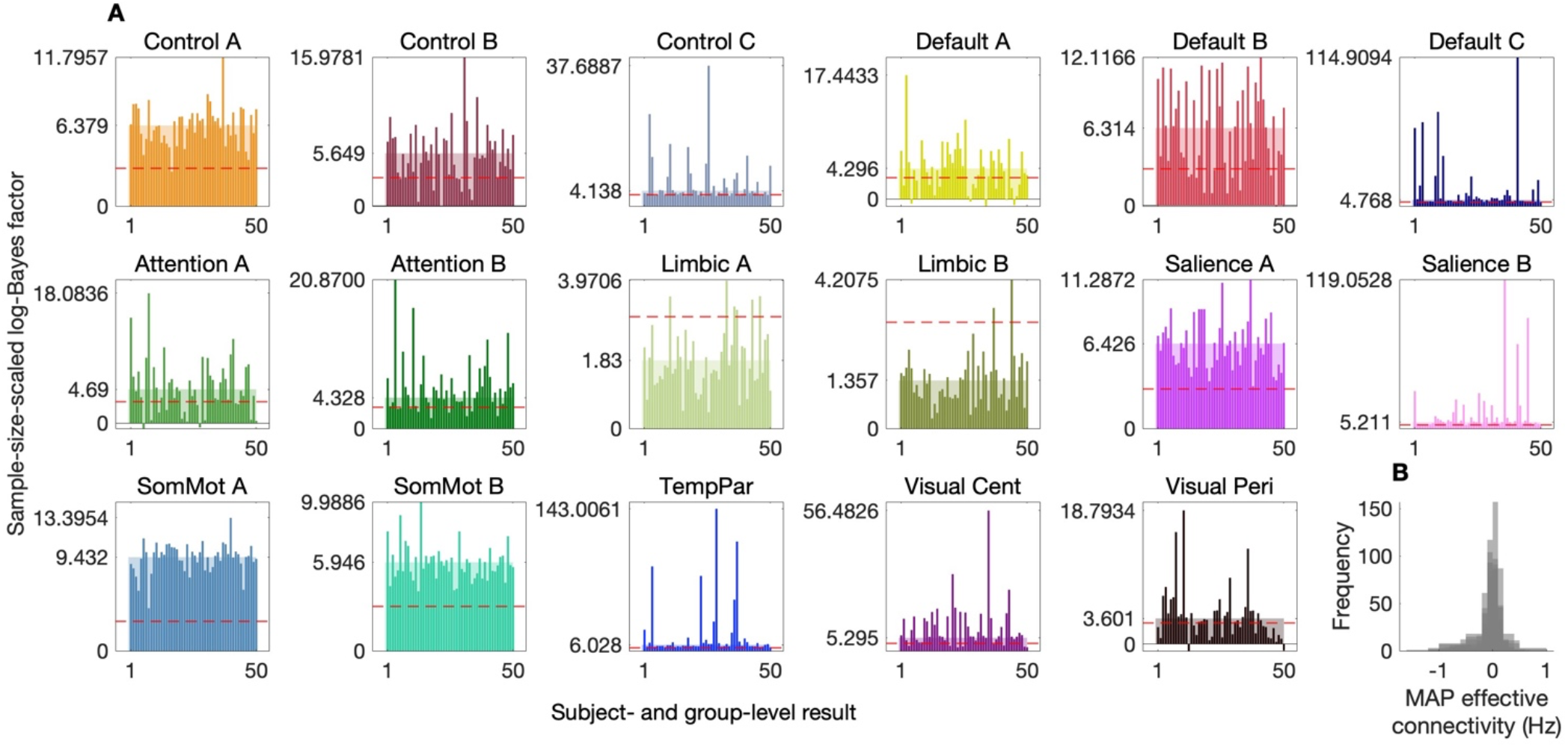
Out-of-sample validation of hierarchical empirical Bayes models. (*A*) Log-Bayes factors comparing hierarchical empirical Bayes models with structure-based priors (derived from out-of-sample evidence-weighted prior-variance transformations) to models with uninformative priors, across 17 effective connectivity brain networks. The semi-transparent bars represent the sample-size-scaled log-Bayes factor for group-level component of the model, while the opaque bars represent the log-Bayes factor for the subject-level component (unaffected by sample-size scaling). The log-Bayes factors indicate substantially greater evidence for models incorporating structure-based priors. Note a dashed red line indicates log-Bayes factor of 3. Nonzero y-axis tick marks indicate the (per-subject) increase in evidence at the group level and the maximum increase in evidence at the subject level, respectively. Networks analyzed include control (Control A–C), default mode (Default A–C), attention (Attention A, B), limbic (Limbic A, B), salience (Salience A, B), somatomotor (SomMot A, B), temporoparietal (TempPar), and visual networks (Visual Cent, central; Visual Peri, peripheral). (*B*) Histogram of maximum *a posteriori* (MAP) group-level effective connectivity estimates for all networks.

### Criterion validity

To explore inter-network differences in the degree to which structural connectivity influences—scales the variance of—effective connectivity, we examined the value of the scale hyperparameter, denoted *β*, that defined the slope of the evidence-weighted prior-variance transformations (Fig. 3A). Fig. 5A shows cortical networks color-coded according to the relative strength of this hyperparameter. Fig. 5B clarifies that structural connectivity had the greatest influence on effective connectivity in the default mode (Default A) network that comprised hub regions: the posterior cingulate and medial prefrontal cortices. Here, we also show that these scale hyperparameters are situated along an axis that describes an approximate unimodal (sensory) to transmodal (integrative) processing hierarchy (17).

**Fig. 5.**
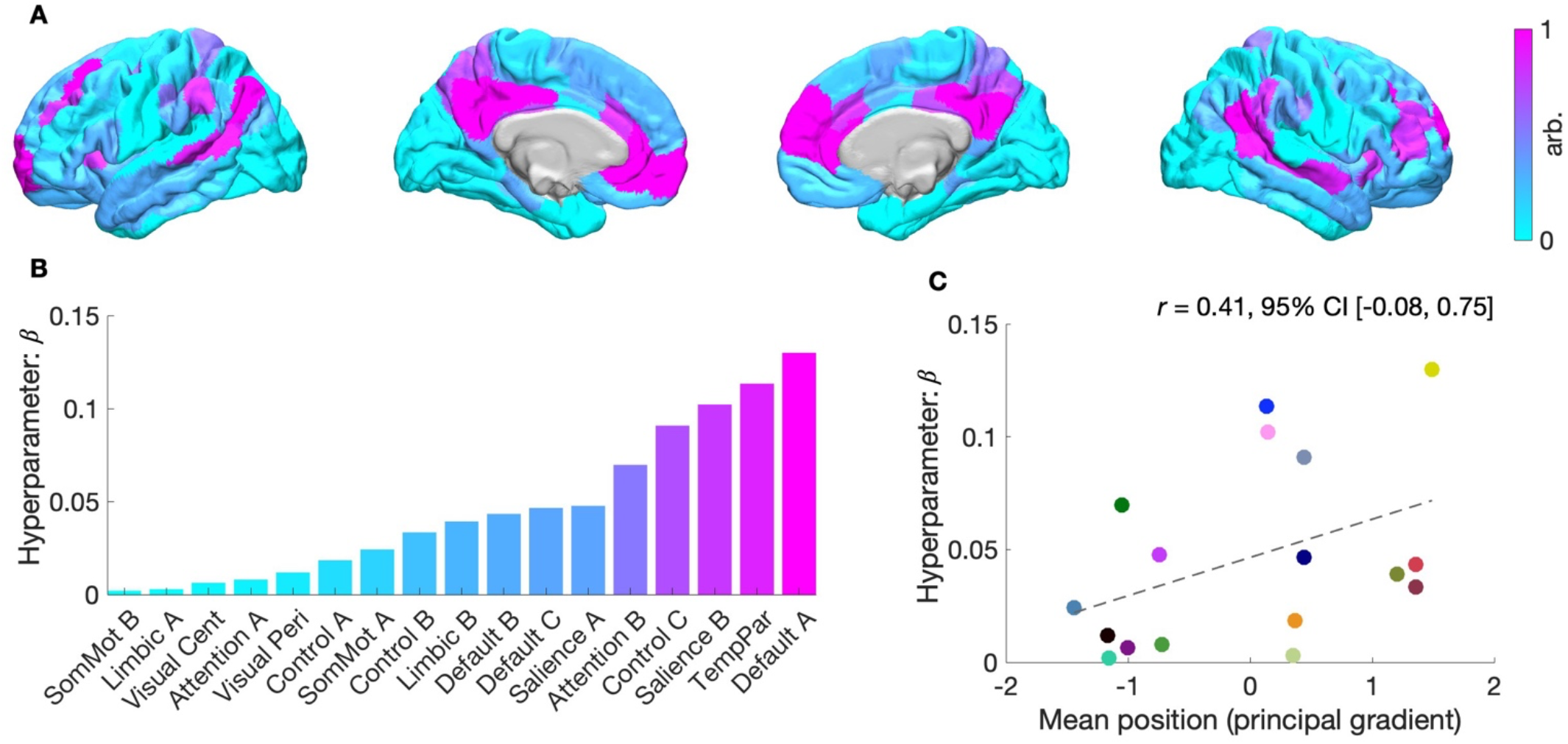
Structural modulation of effective connectivity across large-scale cortical networks. (*A*) Cortical projection of network-specific scale hyperparameters, shown at the parcel level (left lateral, left medial, right medial and right lateral views). Each parcel is colored according to the scale hyperparameter (*β*) of its associated network using a continuous colormap. Higher values (magenta) reflect stronger modulation of effective connectivity by structural connectivity. (*B*) Bar plot of scale hyperparameters ranked by magnitude. The Default A network showed the strongest structural modulation of effective connectivity, while the SomMot B network exhibited the weakest. Networks analyzed include control (Control A–C), default mode (Default A–C), attention (Attention A, B), limbic (Limbic A, B), salience (Salience A, B), somatomotor (SomMot A, B), temporoparietal (TempPar), and visual networks (Visual Cent, central; Visual Peri, peripheral). (*C*) Scatterplot showing the relationship between network-specific scale hyperparameters and the mean position of each network along the principal gradient of functional connectivity (z-scored). The Pearson correlation coefficient (*r*), is reported alongside its 95% confidence interval (square brackets). Scatter colors correspond to the network-specific colors used throughout manuscript.

In this way, notwithstanding that these networks have been examined separately, the results recapitulate a key aspect of a previously identified functional hierarchy (18–20), which in the human cortex, peaks in regions corresponding to the default mode network and reaches its nadir in somatomotor regions (11). Fig. 5C shows a moderate positive correlation (*r* = 0.41, non-significant) between network-specific scale hyperparameters and each network’s mean position along the principal gradient of functional connectivity described by Margulies and colleagues (11). This trend suggests that structural connectivity’s influence on effective connectivity may increase as functional specialization decreases.

## Discussion

This study explored whether integrating structural connectivity into a hierarchical empirical Bayes model improved mapping of resting-state effective connectivity in terms of model evidence. Recently, Sokolov and colleagues introduced a method via which structural connectivity is integrated into a Bayesian RFX model of group-level effective connectivity (7, 8). Here, we built on this prior work in several ways. First, we incorporated BMA into our procedure to account for uncertainty in the selection of prior-variance transformations (rather than focusing on a single best transformation). Second, we examined the impact of utilizing structurally informed group-level effective connectivity as empirical priors for re-evaluating subject-level effective connectivity. Third, we establish the face validity of the hierarchical empirical Bayes model *in silico* and demonstrate that it is more accurate than a structurally informed MVAR model. Fourth, we demonstrated our procedure’s test–retest reliability and out-of-sample validity across resting-state networks. Finally, we showed that inter-network differences in the coupling between structural and effective connectivity—the modulation of prior variance by structural connectivity—recapitulate a well-known cortical hierarchy.

Our findings indicate that structural connectivity constrains resting-state effective connectivity, with the operative *a priori* assumption being that the probability of an interaction between two neuronal populations increases with the extent to which two regions are connected via nerve tracts detectable at the macroscale of MRI. However, this relationship between structural and effective connectivity does not appear to be uniform across the brain but rather appears to be modulated, hierarchically, along an approximate unimodal– transmodal axis (17). The recapitulation of this well-known cortical hierarchy not only attests to the criterion validity of the hierarchical empirical Bayes model, but also hints at a deeper principle of brain organization. Namely, that the principal, unimodal–transmodal gradient of functional connectivity may be explained in terms of the influence that structural connectivity exerts on effective connectivity: an influence that appears to increase with decreasing functional specialization (in other words, as regions become more involved in integrating information across the brain).

While previous studies have shown that structure–function coupling—typically measured via the correlation between each region’s structural and functional connectivity profile (21)—is stronger in unimodal regions and weaker in transmodal regions (18, 22), our findings appear to show the opposite pattern with effective connectivity: we observe a stronger influence of structural connectivity in transmodal networks. This apparent divergence should be interpreted with care, however, given the fundamental differences between functional and effective connectivity. From a function-to-structure (inverse) perspective, one might expect that if source-to-target neuronal communication is more flexible or variable—as is often assumed for transmodal networks (23, 24)—then a broader portion of the structural connectivity space may be sampled by these dynamics. Consequently, even though transmodal regions may appear more ‘untethered’ from anatomy in terms of correlational structure–function coupling, the space of directed interactions (effective connectivity) that best explains observed neural dynamics can nonetheless be more tightly shaped by structural connectivity, consistent with our findings.

Our results are aligned with those from studies that have integrated structural connectivity into effective and directed functional connectivity models via priors. Namely, previous work has shown that introducing a positive, monotonic mapping between structural connectivity and prior variance in the context of dynamic causal modeling and MVAR models, increases model evidence (8, 25–27). More specifically, our work is aligned with prior work examining the impact of structure-based priors in group-level models (8, 28). It differs from this prior work, however, as previous investigations have not, per se, examined the impact of leveraging a structurally informed group-level effective connectivity to constrain subject-level effective connectivity, and have not validated structure-based priors with new, unseen data. Furthermore, rather than utilize a sigmoidal prior-variance transformation per earlier studies (8, 25), we utilized a simpler and more interpretable linear function amenable to analytic BMA (unlike the sigmoid, linear functions are closed under linear combination, meaning the weighted average of linear functions remains linear).

The results of this study are also aligned with those that have involved embedding structural connectivity into MVAR-type models. Recently, Tanner and colleagues demonstrated that structurally informed directed functional connectivity obtained under such a model exhibited a hierarchical community structure (9), and related models have demonstrated predictive validity across several neuropsychiatric conditions (29–31). That said, such models preclude multi-hop directed functional connectivity (directed influences between region *i* to region *j* that are mediated via one or more relay regions). Our *in silico* analyses (Fig. 2E–F; Fig. S1B–C) suggest that this limitation may hinder the ability of these models to accurately characterize directionality (leading to higher sign errors, compared to the hierarchical empirical Bayes approach).

Our study has important implications. First, it underscores the importance of considering structural connectivity in effective connectivity, with findings suggesting that integrating structure-based priors into a hierarchical model of effective connectivity facilitates robust inference. Second, it offers a framework via which one can investigate the coupling between structural and effective connectivity, and how such coupling might be modified. The timeliness of such questions is thrown into sharp contrast considering recent evidence for disorder- and intervention-specific alterations in the hierarchical organization of brain function—with increased coupling between functional and structural connectivity in various neuropsychiatric disorders (21, 32), and psychedelics inducing an uncoupling of functional and structural connectivity (33, 34). Finally, the study demonstrates that the hierarchical empirical Bayes model outperforms a popular alternative approach to characterizing directed influences between brain regions and has yielded findings that hint a mechanism via which the principal, unimodal–transmodal gradient of functional connectivity may emerge. Whether this gradient emerges from the differing constraint that the nerve tracts exert on interregional brain communication is an idea that warrants the attention of future investigations.

This study’s implications need to be considered in view of certain limitations. First, the key limitation to integrating structural connectivity into effective connectivity via structure-based priors is that the integration is somewhat heuristic, representing a statistical assumption rather than a process that maps to a biological mechanism (3). That said, there are myriad ways in which such mechanistic information might be embedded into this hierarchical empirical Bayes model or incorporated into structural connectivity itself by, for example, adding biological annotations (35). Second, owing to the inability of dwMRI to determine axonal directionality (36), our procedure used symmetric structural connectivity, and thus for two given regions, equal priors were assigned for efferent and afferent effective connections. Third, although the use of closed-form solutions (Eqs. 11–12) alongside a linear formulation (Eq. 7) renders the inversion procedure (Fig. 1) highly efficient, we nonetheless inherit the computational constraints of (currently implemented) spectral DCM, which—given realistic resource demands—is best suited to networks comprising up to 64 regions (37), and thus typically precludes whole-brain analyses. Finally, although we made use of two different atlases (SI Results, Fig. S2), both were derived to maximize within-parcel functional homogeneity, without regard for structural connectivity (10, 38). The extent to which such modality-specific parcellations confound measures of structure–function relationships—whether based on simple correlations or structure-based priors—remains an open question (39, 40). Future research can hope to address these issues by exploring graph-theory-based methods of inferring asymmetric signaling from structural connectivity (41, 42), in addition to a wider array of atlases.

In conclusion, our study highlights the important role of structural connectivity in shaping effective connectivity, presenting—to our knowledge—the first evidence of network-dependent modulation of this relationship in humans. Using a novel hierarchical empirical Bayes method, we demonstrate that a positive, monotonic relationship between structural connectivity and the prior variance of effective connectivity generalizes across sessions and samples. The model’s criterion validity is supported by its recapitulation of a well-known cortical hierarchy, while its construct validity is evidenced by superior recovery of subject- and group-level effective connectivity compared to an alternative model. Taken together, these findings recommend a shift towards more integrative approaches in which the fusion of structural and effective connectivity could offer novel insights into functional integration in health and disease.

## Methods

### Data

Data used in this study were sourced from the HCP (15). Details concerning MRI acquisition, preprocessing, tractography, and the derivation of parcellated structural connectivity—as well as the procedures used to generate simulation results (Fig. 2)—are provided in the supporting information (SI Methods).

### Model

The hierarchical empirical Bayes model is used to characterize effective connectivity networks across and within subjects (*s* = 1, …, *S*), comprising *i* = 1, …, *n* neuronal populations. At the subject level, the model builds on the well-known and well-validated DCM for resting-state fMRI (also known as spectral DCM) (4, 43, 44). The temporal-domain formulation assumes the following continuous-time state-space representation:

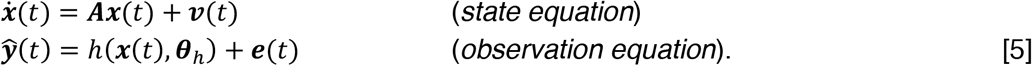

Here, ***x***(*t*) = [*x*_1_(*t*), …, *x*_*n*_(*t*)]^T^ denotes the neuronal state vector, where each scalar function *x*_*i*_(*t*) ∈ ℝ describes the ensemble (or mean-field) activity of the *i*-th population at time *t*. The transition matrix ***A*** ∈ ℝ^*n*×*n*^ encodes the intra- and inter-regional modulation of the rates of change in this ensemble activity (quantifying the effective connectivity), in its diagonal and off-diagonal elements, respectively (with units of Hz, reflecting the continuous-time rate constants). In the observation equation, *h* is the hemodynamic response function (HRF) with free parameters ***θ***_*h*_ (SI Methods), which maps ensemble neuronal activity to expected BOLD responses ***ŷ***(*t*). In spectral DCM, both endogenous fluctuations ***v***(*t*), and observation error ***e***(*t*), are parameterized as power-law noise (SI Methods).

Using the Fourier transform, ℱ, the state-space model is transformed into the spectral domain, such that it generates the expected cross-spectral density (CSD) of BOLD responses ***Ĝ***_*y*_(*ω*) = [ℱ{***ŷ***(*t*)}ℱ{***ŷ***(*t*)}^†^], where † denotes the conjugate transpose, and the right-hand side is implicitly a function of angular frequency *ω* via the Fourier transform. Putting this all together, the spectral equivalent of Eq. 5 reads:

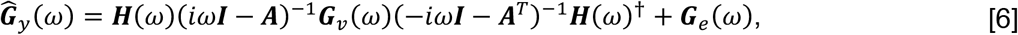

where ***H***(*ω*) is the Fourier transform of the HRF. Here, notably, the latent neuronal state in the frequency domain, ***X***(*ω*), has been factored out via the substitution ***X***(*ω*)***X***(*ω*)^†^ = (*iω****I*** − ***A***)^−1^***G***_*v*_(*ω*)(−*iω****I*** – ***A***^T^)^−1^. To construct the empirical CSD matrix ***G***_*y*_(*ω*) for model inversion, we fit an MVAR model to the data (SI Methods). For a didactic introduction to these details, see Novelli and colleagues (45).

In the equations that follow, we refer to Eq. 6 as the first-, subject-level model 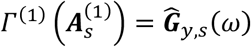, where the subscript *s* denotes the subject-specificity of parameters and (expected) data, and the superscript (1) distinguishes these elements from higher levels of the hierarchy. With the specification of a (multivariate) Gaussian prior over model parameters ***θ***_*s*_ (SI Methods), the statistical model for a given DCM is defined in terms of the joint probability distribution, *p*(***G***_*y,s*_(*ω*), ***θ***_*s*_) = *p*(***G***_*y,s*_(*ω*)|***θ***_*s*_)*p*(***θ***_*s*_). This provides the necessary ingredients for model inversion via VBL (Eq. 2), which approximates the likelihood *p*(***G***_*y,s*_(*ω*)|***θ***_*s*_) under the simplifying assumption that it is Gaussian, reflecting independent and identically distributed (IID) additive Gaussian noise in the data.

With DCMs inverted for a given network, the posteriors of interest—for effective connectivity—are moved into the hierarchical empirical Bayes model (Fig. 1) (14). Vertically stacking the vectorized posterior means for *S* subjects, 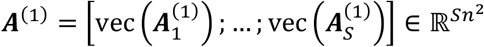, permits the following (vectorized regression) formulation:

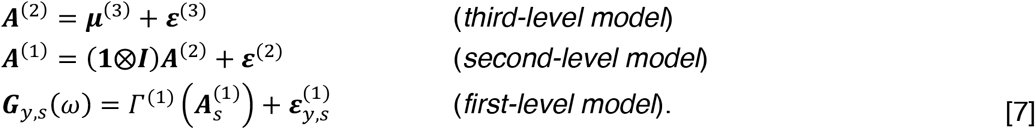

Here, at the third level, 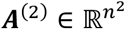 is the group-level effective connectivity modeled as deviations ***ε***^(3)^∼𝒩(**0, Σ**^(3)^) from prior expectations ***μ***^(3)^. At the second level, the Kronecker product **1** 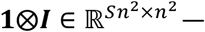 — where **1** ∈ ℝ^*S*^ is a column vector of ones and 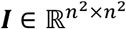 is the identity matrix—ensures that both ***A***^(2)^ and the RFX ***ε***^(2)^, are appropriately tiled to each parameter in ***A***^(1)^. The final line of Eq. 7 models the observed CSD for each subject, ***G***_*y,s*_(*ω*), as the output of the DCM mapping Γ^(1)^ applied to subject-specific effective connectivity 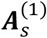—the *s*-th *n*^2^-dimensional sub-vector of ***A***^(1)^—plus residual error 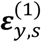.

Following established conventions, ***ε***^(2)^∼𝒩(**0, Σ**^(2)^) is parametrized in terms of a scaled precision component (SI Methods). At the third level, structural connectivity is incorporated into **Σ**^(3)^ via the following:

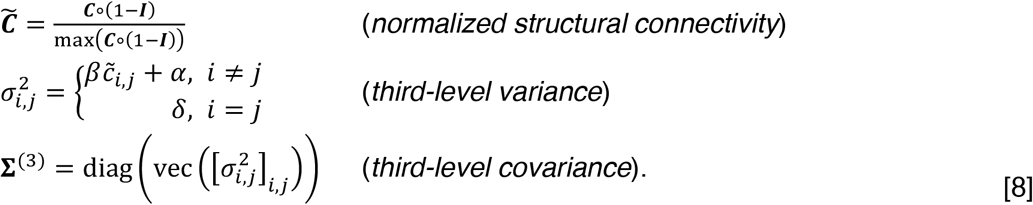

Here, 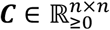 encodes the structural connectivity for an *n*-region network, (1 − ***I***) is the complement of the *n*-dimensional identity matrix, and the normalized structural connectivity weights 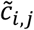 from the *i*-th row and *j*-th column are transformed to produce variance terms 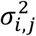 for corresponding group-level effective connections ***A***^(2)^. The hyperparameters *α* and *β* set the baseline variance and its scaling with structural connectivity, respectively, and intraregional connections—where *i* = *j*—are set to *δ*. In the context of our analyses, uninformed models represent the case where *β* = 0, *α* = 1/2 and *δ* = 1/64.

Taken together, the joint distribution over observed data and parameters implied by the hierarchical model can be factorized as:

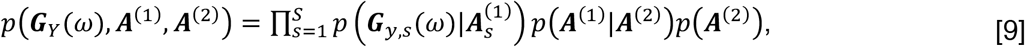

where ***G***_*Y*_(*ω*) = {***G***_*y*,1_(*ω*), …, ***G***_*y,S*_(*ω*)} denotes the observed CSDs for all subjects. The three right-hand components correspond to: first-level likelihoods (of observed CSDs under subject-specific effective connectivity); the second-level model capturing variability across subjects; and the third-level (structure-based) prior over group-level effective connectivity.

Although in principle one could invert this full hierarchical model jointly, in practice we adopt a computationally efficient, multi-step approach utilizing standard tools for Bayesian inference. First, the group-level model *p*(***A***^(1)^, ***A***^(2)^) = *p*(***A***^(1)^|***A***^(2)^)*p*(***A***^(2)^) is inverted using a VBL scheme that incorporates BMR (SI Methods), yielding the posterior *q*(***A***^(2)^|***A***^(1)^) over group-level effective connectivity (first inset, Fig. 1B). Then, we utilize BMR again (second inset, Fig. 1B), exploiting the sufficient statistics of the subject-level priors 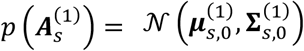 and posteriors 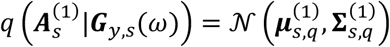, along with the empirical prior furnished by the second-level posterior, 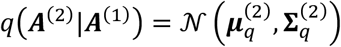, to compute a reduced (updated) posterior 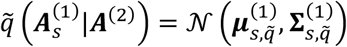, according to:

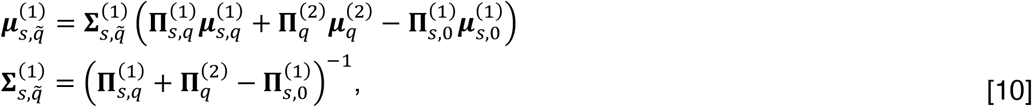

where 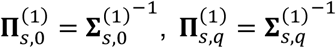, and 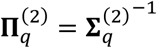. In this way, the second-level posterior propagates downward to inform the update of first-level models, with the update weighted by the precision of subject-level posteriors. The free energy for each reduced first-level model, 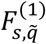, is then computed from the free energy of the original DCM 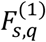, via:

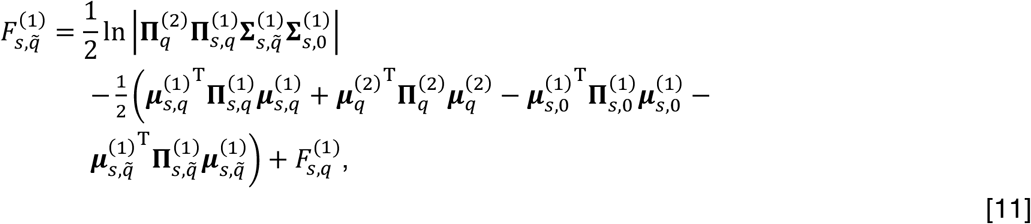

where 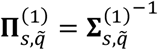. It is important to note that BMR is agnostic to the hierarchical level at which it is applied. This flexibility allows us to compare structurally informed and uninformed models at the group level, propagate the resulting empirical priors downward through the hierarchy, and quantify corresponding changes in model evidence at both the subject and group levels via their respective free energy estimates. This entire BMR procedure is computationally efficient, with all updates performed in closed form.

### Procedures

In our study, for each of the 150 subjects, across 17 networks and two sessions, a spectral DCM was specified and inverted using VBL as implemented in the Statistical Parametric Mapping toolbox (SPM12) (16). We then inverted the hierarchical empirical Bayes model (Steps 1 and 2; Fig. 1) with uninformative third-level priors—*β* = 0, *α* = 1/2 and *δ* = 1/64—for each network within each dataset: the (session-1) test and (session-2) retest datasets (*N* = 100), and two (session-1 and -2) validation datasets (*N* = 50).

In the empirical face-validation phase, we used the test dataset DCMs and BMR to evaluate the evidence associated with different reduced (structurally informed) group-level models, varying the parametrization of the prior-variance transformation (Eq. 8). In these analyses, *α* was sampled at 30 equidistant points across [−1/2, 1/2]. For each *α, β* was sampled at 30 equidistant points across [0, 1/2 − *α*]. Valid parameter combinations, satisfying *α* ≥ 0 and *α* + *β* ≥ 10 ^−5^, were retained, yielding a triangular sampling space with *K* = 7,273 parameter regimes. For each network, we then weighted the *k*-th set of third-level variance parameters, 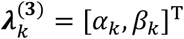, by their corresponding posterior model probabilities 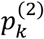, computed via a softmax transformation of the model free energies 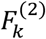. Using BMA, the parameters of the evidence-weighted prior-variance transformation were then obtained as:

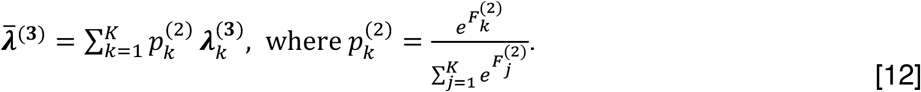

This procedure serves to identify the parameters of a parsimonious transformation (avoiding overfitting) and to quantify uncertainty around those parameters (46) (Figs. 2–3). The resulting parameters 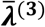, can thus be interpreted as a conservative solution that retains the influence of plausible alternative transformations.

The network-specific evidence-weighted prior-variance transformations identified in the test dataset were applied out of session (in the retest dataset) and out of sample (in the validation datasets). This testing proceeded as follows. After inverting network-specific DCMs for these datasets, we inverted a hierarchical empirical Bayes model—per network—under uninformative priors (using VBL for the group-level model and BMR to update subject-level models). We then used the evidence-weighted prior-variance transformations to construct structurally informed priors and re-inverted each hierarchical model *de novo*. This yielded updated posteriors and free energies at both the group and subject levels. The free energy of these structurally informed models was then compared to that of the uninformed models, at the group level (semi-transparent bars, Fig. 4) and subject level (opaque bars, Fig. 4).

### Data and materials availability

Empirical MRI data analyzed in this study are publicly available, subject to Human Connectome Project (HCP) registration and data-use terms. The code and derived data underlying our implementation of the hierarchical empirical Bayes model—including the *in silico* analyses, permutation-based control analyses, and materials needed to reproduce the results figures—are publicly available on GitHub at: https://github.com/mdgreaves/hierarchical-empirical-Bayes

## Acknowledgments

Data were provided by the Human Connectome Project, WU-Minn Consortium (Principal Investigators: David Van Essen and Kamil Ugurbil; 1U54MH091657) funded by the 16 NIH Institutes and Centers that support the NIH Blueprint for Neuroscience Research; and by the McDonnell Center for Systems Neuroscience at Washington University. M.D.G. was supported by an Australian Government Research Training Program Scholarship. M.D.G., L.N. and A.R. were funded by the Australian Research Council (ref. DP200100757). A.R. was also funded by Australian National Health and Medical Research Council Investigator Grant (ref. 1194910). A.R. was affiliated with The Wellcome Centre for Human Neuroimaging supported by core funding from Wellcome (203147/Z/16/Z). A.R. was a CIFAR Azrieli Global Scholar in the Brain, Mind & Consciousness Programme.

## Author contributions

Conceptualization: M.D.G, L.N., and A.R. Methodology: M.D.G, L.N., A.R. Investigation: M.D.G. Visualization: M.D.G. Supervision: L.N. and A.R. Writing—original draft: M.D.G. Writing—review & editing: M.D.G., L.N., and A.R.

