## Supporting Information for "Structurally informed resting-state effective connectivity recapitulates cortical hierarchy"

### Overview
This supporting information (SI) document provides additional analyses, figures, and methodological details that support the findings reported in the main text. Elements appear in the order in which they are referenced in the main text. A complete list of contents is provided in the table below for ease of navigation.

### SI Methods

#### Completing the hierarchical empirical Bayes model

To complete the specification of the hierarchical empirical Bayes model (Methods, Eqs. 5–8), we detail the form of the HRF, endogenous fluctuations, observation error, the construction of the empirical CSD matrix, the parametrization of residual error (in the context of VBL) and the second-level prior covariance for RFX. In Eq. 5 (Methods), $h(\boldsymbol{x}(t), \boldsymbol{\theta}_h)$ represents the well-known 'Balloon' HRF that yields expected BOLD responses from regional neuronal states $x_i(t)$ as follows (1):

$$
\begin{aligned}
\dot{s}_i(t) &= x_i(t) - k_h \cdot s_i(t) - \gamma_h \cdot (f_i(t) - 1), && (\textit{vasodilatory signal}) \\
\dot{f}_i(t) &= s_i(t), && (\textit{blood flow induction}) \\
\dot{b}_i(t) &= \frac{1}{\tau_i} \cdot \left( f_i(t) - b_i(t)^{1/\alpha_h} \right), && (\textit{blood volume}) \\
\dot{q}_i(t) &= \frac{1}{\tau_i} \cdot \left( \frac{f_i(t) \cdot \left( 1 - (1 - E_0)^{1/f_i(t)} \right)}{E_0} - q_i(t) \cdot b_i(t)^{1/\alpha_h - 1} \right), && (\textit{deoxyhemoglobin content}) \\
\hat{y}_i(t) &= V_0 \cdot \left[ k_1 \big(1 - q_i(t)\big) + k_2 \left(1 - \frac{q_i(t)}{b_i(t)}\right) + k_3 \big(1 - b_i(t)\big) \right]. && (\textit{BOLD responses})
\end{aligned}
\qquad [S1]
$$

Here, neuronal activity $x_i(t)$ drives a vasodilatory signal $s_i(t)$, which increases blood flow $f_i(t)$, expands venous blood volume $b_i(t)$ via Grubb's law, alters deoxyhemoglobin content $q_i(t)$ through oxygen extraction, and together these variables determine the expected BOLD response $\hat{y}_i(t)$. The vasodilatory signal decays at rate $k_h$, the feedback of blood flow is regulated by $\gamma_h = 0.32$, $\boldsymbol{\tau} = [\tau_1, \dots, \tau_n]$ are region-specific transit times, $\alpha_h = 0.32$ is Grubb's (vessel stiffness) exponent, and $E_0 = 0.4$ and $V_0 = 4$ represent the resting oxygen extraction fraction and resting venous blood volume fraction, respectively. In the last line of Eq. S1, coefficients $k_1$, $k_2$, and $k_3$ represent the contributions of the intra- and extra-vascular compartments to the BOLD response:

$$
k_1 = 4.3 \cdot v_0 \cdot E_0 \cdot TE, \qquad k_2 = \epsilon_h \cdot r_0 \cdot E_0 \cdot TE, \qquad k_3 = 1 - \epsilon_h, \qquad [S2]
$$

where $v_0 = 40.3$ is the frequency offset at the outer surface of magnetized vessels, $TE = 0.04$ is the echo time, $r_0 = 25$ is the slope of the intravascular relaxation rate, and $\epsilon_h$ is the ratio of intra- to extra-vascular signal contributions. The set of parameters that were free to vary is given by $\boldsymbol{\theta}_h = \{k_h, \boldsymbol{\tau}, \epsilon_h\}$.

We now turn to the model of endogenous fluctuations $\boldsymbol{G}_v(\omega) \in \mathbb{R}^{n \times n}$, and observation error $\boldsymbol{G}_e(\omega) \in \mathbb{R}^{n \times n}$ (Methods, Eq. 6). Both models take the form of a diagonal matrix-valued function of angular frequency $\omega$, whose entries—$g_{v,i,i}(\omega)$ and $g_{e,i,i}(\omega)$, respectively—encode an auto-spectrum or power spectral density (PSD) model of region-specific endogenous fluctuations and observation error:

$$
g_{v,i,i}(\omega) = \alpha_v \omega^{-\beta_v}, \qquad g_{e,i,i}(\omega) = \big(\gamma_e \cdot \alpha_{e,i}\big) \cdot \frac{\omega^{-\beta_e/2}}{\sum_\omega \omega^{-\beta_e/2}}. \qquad [S3]
$$

Here, the PSD model for endogenous fluctuations is parameterized by a global amplitude $\alpha_v$ and spectral exponent $\beta_{v,i}$, whereas the PSD model for observation error is controlled by a global spectral exponent $\beta_e$, and region-specific amplitude parameter $\alpha_{e,i}$ normalized by a global factor $\gamma_e$.

To construct the empirical CSD matrix $\boldsymbol{G}_y(\omega)$ for model inversion, we fit a multivariate autoregressive model of order $p = 8$ to observed BOLD time series using the variational Bayesian procedure described by Penny and Roberts (2). This procedure yields a set of autoregressive coefficient matrices $\{\boldsymbol{W}_k \in \mathbb{R}^{n \times n}\}_{k=1}^{p}$ and a residual covariance matrix $\boldsymbol{\Sigma}_w \in \mathbb{R}^{n \times n}$. The standard spectral factorization then produces $\boldsymbol{G}_y(\omega)$ as:

$$\boldsymbol{G}_y(\omega) = \boldsymbol{\mathcal{H}}(\omega)\boldsymbol{\Sigma}_w\boldsymbol{\mathcal{H}}(\omega)^\dagger, \text{ where } \boldsymbol{\mathcal{H}}(\omega) = \left(\boldsymbol{I} + \sum_{k=1}^{p} \boldsymbol{W}_k\, e^{-i\omega k}\right)^{-1}, \quad \text{[S4]}$$

Consistent with prior related work, $\boldsymbol{G}_y(\omega)$ is estimated at $n_f = 32$ linearly spaced frequencies between $1/128$ Hz and the Nyquist frequency: $1/2 \cdot TR$, where $TR$ is the fMRI repetition time (sampling interval) (3).

In the context of model inversion, the expected log-likelihood $\mathbb{E}_{q(\boldsymbol{\theta})}[\ln p(\boldsymbol{y}|\boldsymbol{\theta})]$ (Results, Eq. 2), quantifies the fit between the expected and observed CSDs under the approximate posterior. This is expressed as:

$$\text{vec}\left(\boldsymbol{G}_y(\omega)\right) = \text{vec}\left(\widehat{\boldsymbol{G}}_y(\omega)\right) + \boldsymbol{\epsilon}_y. \quad \text{[S5]}$$

In the context of VBL, the residual error $\boldsymbol{\epsilon}_y \sim \mathcal{N}(\boldsymbol{0}, \boldsymbol{\Sigma}_y)$ is parameterized in terms of its precision. Per the implementation utilized here, the residual error precision $\boldsymbol{\Pi}_y = \boldsymbol{\Sigma}_y^{-\boldsymbol{1}}$, is expressed as a weighted sum of $n^2$ precision components:

$$\boldsymbol{\Pi}_y = \sum_{i=1}^{n^2} e^{\lambda_{y,i}}\, \boldsymbol{Q}_{y,i}, \quad \text{[S6]}$$

where $\boldsymbol{Q}_{y,i} \in \mathbb{R}^{(n^2 \cdot n_f)\times(n^2 \cdot n)}$ is a sparse, block-diagonal matrix with an $n_f$-dimensional identity sub-block targeting the $i$-th region pair (including self-connections). Here, the precision weights $\lambda_{y,i}$ are drawn from a hyperprior $\lambda_{y,i} \sim \mathcal{N}(8, 1/128)$, where these settings were determined elsewhere, *in silico* (4).

Finally, in the group-level model (Methods, Eq. 7), $\boldsymbol{\varepsilon}^{(2)} \sim \mathcal{N}(\boldsymbol{0}, \boldsymbol{\Sigma}^{(2)})$ are parametrized in terms of a scaled precision component:

$$\boldsymbol{\Sigma}^{(2)^{-1}} = \boldsymbol{\Pi}^{(2)} = \boldsymbol{I} \otimes \left(\boldsymbol{Q}_0^{(2)} + e^{-\gamma_q^{(2)}} \boldsymbol{Q}_1^{(2)}\right), \quad \text{[S7]}$$

where $\boldsymbol{I}$ is an $S$-dimensional identity matrix, $\boldsymbol{Q}_0^{(2)} = e^{-8}\boldsymbol{\Sigma}^{(3)^{-1}}$ is the lower-bound precision, $\boldsymbol{Q}_1^{(2)} = 16 \cdot \boldsymbol{\Sigma}^{(3)^{-1}}$, and the precision scale parameter $\gamma_q^{(2)}$ is inferred from the data (Table S1).

| Parameter | Description | Prior mean | Prior variance |
|---|---|---|---|
| $\ln\left(-2\cdot a_{s,i,i}^{(1)}\right)$ | First-level intra-regional effective connectivity | $-1/2$ | $1/64$ |
| $a_{s,i,j}^{(1)}$ | First-level inter-regional effective connectivity | $1/128$ | $1/2$ |
| $\ln\alpha_v$ | Global amplitude of endogenous fluctuations | $0$ | $1/64$ |
| $\ln\beta_v$ | Global spectral exponent of endogenous fluctuations | $0$ | $1/64$ |
| $\ln\alpha_{e,i}$ | Regional amplitude of observation noise | $0$ | $1/64$ |
| $\ln\beta_e$ | Global spectral exponent of observation error | $0$ | $1/64$ |
| $\ln\gamma_e$ | Normalization factor for observation error | $0$ | $1/64$ |
| $\ln k_h$ | Decay rate of the vasodilatory signal | $0$ | $1/256$ |
| $\ln\epsilon_h$ | Ratio of intra- to extra-vascular signal contributions | $0$ | $1/256$ |
| $\ln\tau_i$ | Transit times | $0$ | $1/256$ |
| $\ln\left(-2\cdot a_{i,i}^{(2)}\right)$ | Second-level intra-regional effective connectivity | $-1/2$ | $1/64$ |
| $a_{i,j}^{(2)}$ | Second-level inter-regional effective connectivity | $1/128$ | Eq. 8 |
| $\gamma_q^{(2)}$ | Second-level precision scale parameter | $0$ | $1/16$ |

**Table S1.** Prior mean and variance for free parameters utilized in the hierarchical empirical Bayes model. Notation is consistent with that utilized in the main text. If the model is structurally informed, the second-level inter-regional effective connectivity is a function of normalized structural connectivity, and the hyperparameters governing the prior-variance transformation (Methods, Eq. 8).

### Conducting *in silico* analyses

To evaluate the face validity of the hierarchical empirical Bayes model, we conducted a test of model identifiability (parameter recovery), and construct validation in which we compared the model to a structurally informed MVAR model. These analyses involved the simulation of group- and subject-level effective connectivity, neuronal dynamics, and BOLD signals making use of standard procedures implemented via the SPM toolbox (version 12) (4, 5).

First, we obtained a random, sparse, symmetric matrix $\boldsymbol{C}\in\mathbb{R}_{\geq 0}^{n\times n}$ with density $d_c$. Here, in these analyses, this group-level structural connectivity $\boldsymbol{C}$, satisfied the condition that at least one region was completely unconnected; that is, there existed at least one row (column) that is entirely zero (Results, Fig. 2A). After normalization (Methods, Eq. 8), we sampled random variates for $\boldsymbol{A}^{(2)}$ from its prior (Table S1), under fixed hyperparameter values (Table S2). We then obtained subject-level effective connectivity parameters $\boldsymbol{A}_s^{(1)}$, for $s=1,\dots,S$, by perturbing $\boldsymbol{A}^{(2)}$ with randomly sampled noise $\boldsymbol{\varepsilon}^{(2)}\sim\mathcal{N}(\boldsymbol{0},\boldsymbol{\Sigma}^{(2)})$, where $\boldsymbol{\Sigma}^{(2)}=\delta\boldsymbol{I}$, and $\boldsymbol{I}\in\mathbb{R}^{Sn^2\times Sn^2}$ is the identity matrix.

| Parameter | Description | Parameter value |
|---|---|---|
| $S$ | Number of instantiations (simulated subjects) | $50$ |
| $n$ | Number of regions | $6$ |
| $T$ | Number of time points (scans) | $1000$ |
| $TR$ | Repetition time | $1$ |
| $\mathrm{SNR}$ | Signal-to-noise ratio (linear) | $\{1,5,10,50,100,200\}$ |
| $d_c$ | Structural connectivity density | $2/3$ |
| $\alpha$ | Baseline variance of group-level effective connectivity $a_{i,j}^{(2)}$ | $1/16$ |
| $\beta$ | Influence of structural connectivity on variance of $a_{i,j}^{(2)}$ | $1/10$ |
| $\delta$ | Variance of subject-level effective connectivity | $1/64$ |
| $k_h$ | Decay rate of the vasodilatory signal | $1$ |
| $\epsilon_h$ | Ratio of intra- to extra-vascular signal contributions | $1$ |
| $\tau_i$ | Transit times | $\sim \mathrm{LogNormal}(0, e^{-8})$ |

**Table S2.** Hardcoded parameter values used in simulations. Notation is consistent with that utilized in the main text. For each simulation of $S$ instantiations (simulated subjects), the hemodynamic transit times $\tau_i$ were sampled independently for each region ($i = 1, \dots, n$) from a log-normal distribution.

Next, endogenous fluctuations were simulated as independent first-order autoregressive processes with autoregressive coefficient 0.5, scaled to a stationary standard deviation of 0.25; observation noise was generated similarly, with variance determined by the desired signal-to-noise ratio (SNR). Synthetic BOLD signals were then obtained by integrating the (time-domain) generative model (Methods, Eq. 5).

Finally, we inverted the hierarchical empirical Bayes model (Methods), and additionally estimated directed functional connectivity using a structurally informed MVAR model fitted via ordinary least squares:

$$\boldsymbol{y}[k] = (\boldsymbol{W} \odot \boldsymbol{C}^*)\boldsymbol{y}[k-1] + \boldsymbol{\eta}, \qquad \boldsymbol{C}^* = \left[\widetilde{\boldsymbol{C}} > 0\right], \tag{S8}$$

Here, $k = 1, \dots, T$ indexes discrete time points, where $T$ is the total number of scans; $\boldsymbol{y}[k] \in \mathbb{R}^n$ denotes BOLD activity, $\boldsymbol{W} \in \mathbb{R}^{n \times n}$ is the weight matrix capturing the degree to which the past activity of one region predicts the future activity of another, $\odot$ denotes elementwise multiplication, $\boldsymbol{C}^* \in \{0,1\}^{n \times n}$ is a binarized structural mask, and $\boldsymbol{\eta} \in \mathbb{R}^n$ is a bias term (6). Here, Iverson bracket notation is used $\left[\widetilde{\boldsymbol{C}} > 0\right]$ to indicate elementwise binarization: non-zero elements are set to one.

To evaluate model performance, we used the RMSE, and a macro F1-score computed across three effective connection classes: positive, negative, and absent. Per Razi and colleagues (4), RMSE quantified the average deviation between estimated and ground-truth effective connectivity, while a standard macro F1-score—see Sokolova and Lapalme for didactic introduction to macro-averaging (7)—captured classification performance across all classes. To define class labels, we applied a small threshold ($\zeta = 10^{-5}$) such that values greater than $\zeta$ were classified as positive, less than $-\zeta$ as negative, and within $[-\zeta, \zeta]$ as absent.

#### Second-level inversion, free energy, and scaling of log-Bayes factors

At the second level of the hierarchical empirical Bayes model, inference is performed by maximizing a VBL approximation to the group-level log-model evidence (free energy). Crucially, this objective is not constructed from subject-level posterior means alone: rather, for a given candidate second-level model, BMR is applied subject-wise to evaluate reduced first-level free energies under the implied empirical prior, and these terms are then summed to furnish the fixed-effects 'accuracy' component of the objective that is optimized by iterative ascent. Using subscripts 0 and $q$ to denote prior and approximate posterior quantities, respectively, the second-level free energy is:

$$F_{m,1}^{(2)} = \sum_{s=1}^{S}\left(F_{s,\tilde{q}}^{(1)}\left(\boldsymbol{A}^{(2)}, \gamma_q^{(2)}\right)\right) - \left[\frac{1}{2}\left(\boldsymbol{\mu}_q^{(2)} - \boldsymbol{\mu}_0^{(3)}\right)^{\mathrm{T}} \boldsymbol{\Pi}_0^{(3)}\left(\boldsymbol{\mu}_q^{(2)} - \boldsymbol{\mu}_0^{(3)}\right) + \frac{1}{2}\left(\boldsymbol{\mu}_{q,\gamma}^{(2)} - \boldsymbol{\mu}_{0,\gamma}^{(2)}\right)^{\mathrm{T}} \boldsymbol{\Pi}_{0,\gamma}^{(2)}\left(\boldsymbol{\mu}_{q,\gamma}^{(2)} - \boldsymbol{\mu}_{0,\gamma}^{(2)}\right) + \frac{1}{2}\ln\left|\boldsymbol{C}_0^{-1}\boldsymbol{C}_q\right|\right], \qquad \text{[S9]}$$

where the subtracted 'complexity' term corresponds to the Kullback–Leibler divergence between the joint second-level posterior and its corresponding prior. Here, $F_{s,\tilde{q}}^{(1)}$ is the free energy for the $s$-th reduced first-level model under the empirical prior $q(\boldsymbol{A}^{(2)}|\boldsymbol{A}^{(1)}) = \mathcal{N}\left(\boldsymbol{\mu}_q^{(2)}, \boldsymbol{\Sigma}_q^{(2)}\right)$ (Methods, Eq. 11), whose posterior covariance $\boldsymbol{\Sigma}_q^{(2)}$ depends on the scale of second-level error precision $q(\boldsymbol{\gamma}^{(2)}) = \mathcal{N}\left(\boldsymbol{\mu}_{q,\gamma}^{(2)}, \boldsymbol{\Sigma}_{q,\gamma}^{(2)}\right)$. The quantities $\boldsymbol{\mu}_0^{(3)}$ and $\boldsymbol{\Pi}_0^{(3)}$ are the third-level prior mean and precision for the group-level effective connectivity, respectively (Methods, Eq. 8), while $\boldsymbol{\mu}_{0,\gamma}^{(2)}$ and $\boldsymbol{\Pi}_{0,\gamma}^{(2)}$ are the prior means and precision for the second-level error-precision scale. Finally, $\boldsymbol{C}_0^{-1} = \mathrm{blockdiag}(\boldsymbol{\Pi}_0^{(3)}, \boldsymbol{\Pi}_{0,\gamma}^{(2)})$ is the joint prior precision over $\boldsymbol{A}^{(2)}$ and $\gamma_q^{(2)}$, and $\boldsymbol{C}_q$ is the corresponding posterior covariance.

Because the accuracy component accumulates additively across subjects, differences in second-level free energy—and hence second-level log–Bayes factors comparing alternative hierarchical models—will generally increase in magnitude with sample size, even when the per-subject contribution is modest. To facilitate interpretation in relation to well-known Bayes factor heuristics (8), we report second-level log–Bayes factors scaled by sample size, obtained by dividing the group-level log–Bayes factor by the number of subjects (9).

### Data

Data used in this study were sourced from the HCP (10). Test (face-validation) and retest datasets included data from 100 healthy adults (54 female, age 22–35) acquired on two different days (session 1 and session 2, respectively)*; and two out-of-sample validation datasets included data from 50 healthy adults (24 female, age 22–35) acquired on two different days†.

#### ***Data acquisition and pre-processing***

Data comprised both dwMRI and resting-state fMRI scans from each participant. The dwMRI data were obtained using a spin-echo echo-planar imaging (EPI) sequence constrained by a repetition time (TR) of 5,520 ms, echo time (TE) of 89.5 ms, a flip angle of 78 degrees, and a multiband factor of 3. The field of view (FOV) was set to 210 mm in the readout (RO) direction and 180 mm in the phase encoding (PE) direction, with a resolution matrix of 168 x 144 (RO x PE). Per subject, the data consisted of 111 slices with a 1.25 mm isotropic voxel size, and diffusion-weighted measurements were performed with b-values of 1,000, 2,000, and 3,000 s/mm2, and six b0 scans were also acquired for signal normalization.

*Subject identifiers: 100206, 100307, 100408, 100610, 101006, 101107, 101309, 101915, 102109, 102311, 102513, 102614, 102715, 102816, 103010, 103111, 103212, 103414, 103515, 103818, 104012, 104416, 104820, 105014, 105115, 105216, 105620, 105923, 106016, 106319, 106521, 106824, 107018, 107321, 107422, 107725, 108020, 108121, 108222, 108323, 108828, 109123, 109830, 111009, 111211, 111312, 111413, 111514, 111716, 112112, 112314, 112516, 112920, 113215, 113316, 113619, 113922, 114217, 114419, 114621, 114823, 115017, 115320, 115724, 115825, 116221, 116524, 116726, 117021, 117122, 117324, 117930, 118124, 118225, 118528, 118730, 118831, 118932, 119025, 119732, 119833, 120111, 120212, 120414, 120515, 120717, 121416, 122317, 122418, 122620, 122822, 123117, 123420, 123521, 123723, 123925, 124624, 124826, 125222, 125424.

†Subject identifiers: 108525, 110007, 110411, 110613, 119126, 121618, 121921, 124220, 124422, 125525, 126325, 129634, 129937, 130013, 130114, 130316, 130417, 130720, 131217, 131419, 131823, 131924, 132017, 133019, 133625, 133928, 134021, 134223, 134627, 134728, 134829, 135124, 135225, 135528, 135629, 135730, 135932, 136126, 136227, 136631, 137532, 137633, 138130, 138231, 138332, 139839, 140117, 140319, 140420, 140824.

The fMRI data were acquired using a gradient-echo EPI sequence constrained by a TR of 720 ms, a TE of 33.1 ms, and a flip angle of 52 degrees, and employed a multiband factor of 8. The FOV for rfMRI data was 208 x 180 mm (RO x PE), with a resolution matrix of 104 x 90 (RO x PE). Per participant, the data consisted of 72 slices with an isotropic voxel size of 2 mm. The entire resting-state fMRI scanning session lasted for 14 m 33 s, yielding 1,200 time points of functional brain activity-related data. The HCP protocol includes data acquired using two phase-encoding directions: left-to-right (L–R) and right-to-left (R–L). For our study, we used the L–R phase direction in each dataset.

The HCP minimally pre-processed pipeline (version 3.19.0) was employed for pre-processing the data, with denoising of the functional timeseries data completed using the independent component analysis-based X-noiseifier (FIX) algorithm (11). We refer the reader to the extensive HCP documentation for further details regarding these acquisition and pre-processing steps (12).

#### *Tractography*

Deterministic tractography was performed utilizing the fiber assignment by continuous tractography (FACT) algorithm via MRtrix3 (13, 14). First, at each voxel, a single fiber orientation was modelled by the primary eigenvector of a diffusion tensor fitted to DWI data using iterative least squares. Second, 10 million streamlines were propagated voxel-wise, following the direction of the most aligned (colinear) fiber orientations. Streamlines were dynamically seeded from regions in which streamlines were sparse (relative to the fiber density suggested by the underlying fiber orientation distributions) (15), and either terminated at the gray matter-white matter interface, or at a point where fractional anisotropy (FA) fell below 0.2, curvature exceeded 45 degrees, or length exceeded 250 mm. Finally, these streamlines were downsampled by a factor of 5, and anatomically constrained tractography (ACT) was used to improve their biological plausibility (16).

#### *Network parcellation*

In our main analyses, fMRI and tractography data were parcellated into distinct regions of interest using the 200-region cortical atlas developed by Schaefer and colleagues (17). This atlas, which was generated based on a gradient-weighted Markov random field model and resting-state functional connectivity, can be organized according to 17 resting-state networks identified by Yeo and colleagues (18). Each network, on average, consists of approximately 12 distinct regions, which permitted us to effectively investigate network-level dynamics across the cortex while managing the computational complexity associated with inverting large network models.

Tractography data were parcellated using MRtrix3 and a radial search procedure in which streamlines were assigned to parcels within a 5-mm radius of their endpoints. The fMRI data were parcellated—averaged—using standard HCP workbench command tools. Parcellation yielded (200 x 200) structural connectivity matrices, and (200 x 1,200) fMRI data matrices (capturing time-varying activity across 200 brain regions over 1,200 time points).

With regards to network nomenclature, the 17 networks in Schaefer and colleagues' atlas include: control (Control A–C), default mode (Default A–C), attention (Attention A, B), limbic (Limbic A, B), salience (Salience A, B), somatomotor (SomMot A, B), temporoparietal (TempPar), and visual networks (Visual Cent, central; Visual Peri, peripheral). It is important to note that these network labels are heuristic in nature: they should be considered as providing a useful but simplified representation of the functional roles that these networks play (18). It is also important to note that although some network labels overlap with those in Gordon and colleagues atlas (used in supplementary analyses) (19), the actual network topographies share minimal correspondence (20).

#### *Structural connectivity*

Tractography exhibits well-documented biases whereby short-range connections are overrepresented, overshadowing longer-distance tracts; and larger brain regions appear disproportionately connected (connectivity is positively correlated with regional surface area) (21). To account for these issues, for both the test–retest and validation samples, we generated distance-based consensus structural connectivity matrices per the approach described by Betzel and colleagues (22). These matrices retained the distributions of within-hemisphere and between-hemisphere connection lengths (as approximated via the Euclidean distances between parcel centroids). Connections were divided into 32 distance bins, and edges were selected for inclusion in the group matrix based on their consistency across subjects within each bin. The resultant unweighted consensus matrix was multiplied, elementwise, with an average structural connectivity matrix (the mean connectivity values between all pairs of regions across subjects' data). The resultant weighted consensus matrix was normalized by parcel surface area, yielding a group-level connectome of normalized connection weights.

#### *Cortical gradient of functional connectivity*

Margulies and colleagues applied a diffusion embedding algorithm to fMRI data to extract latent components, referred to as gradients, underlying functional connectivity (23). The principal gradient separated unimodal and transmodal regions along an approximate sensory–fugal axis (24), with somatomotor and default mode regions situated at opposite ends. Leveraging this principal gradient as a quantitative description of this spectrum, we downsampled the gradient from vertex to network resolution by averaging gradient values per each of the 17 Schaefer networks and compared these mean principal gradient values to the scale hyperparameters associated with network-specific evidence-weighted prior-variance transformations.

### SI Results

#### Classification of effective connections *in silico*

Here, Fig. S1A shows the RMSE assessed across the simulated subjects (SNR of 1). Although both the MVAR and hierarchical empirical Bayes model tended to exhibit relatively low error, the latter model demonstrated a tendency toward increased error in certain instances. In contrast, the MVAR model yielded more stable RMSE values across subjects. Fig. S1B shows the polarity error (the percentage of sign errors) assessed across the simulated subjects (SNR of 1), with results indicating that the MVAR model consistently struggled to correctly capture the directionality of connections. Finally, when assessed using the macro F1-score across varying SNRs (Fig. S1C), the group-level estimates obtained under the hierarchical empirical Bayes model consistently outperformed those obtained under the MVAR model.

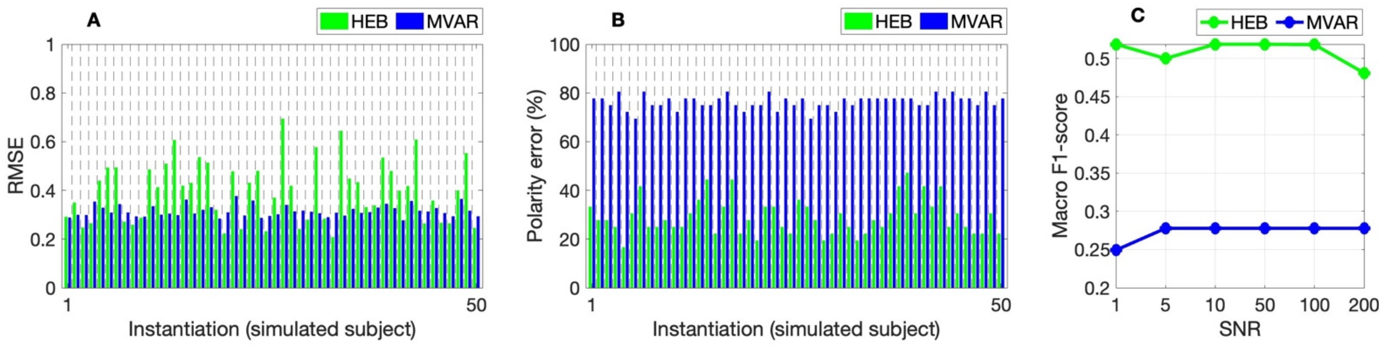

**Fig. S1.** Additional results from *in silico* evaluation of hierarchical empirical Bayes model. (*A*) Bar plot of root mean squared error (RMSE) comparing true and estimated subject-level effective connectivity characterized via the hierarchical empirical Bayes (HEB) model and multivariate autoregressive (MVAR) model across 50 instantiations, highlighting the more stable RMSE values for the latter model. (*B*) Bar plot of polarity error (the percentage of sign errors) comparing the true and estimated subject-level effective connectivity characterized via the HEB model and MVAR model across 50 instantiations, highlighting the greater performance of the former model. (*C*) Macro F1-scores demonstrate that the HEB model more accurately classifies positive, negative, and absent group-level effective connections across varying signal-to-noise ratio (SNR) levels.

### Robustness to atlas selection

To assess the robustness of our results to parcellation, we repeated our exploratory analyses using Gordon and colleagues' (333-region, 13-network) atlas (19), making use of the retest dataset. Several Gordon networks are large, posing challenges for dynamic causal modelling (25). To address this, we applied a clustering-based dimensionality reduction procedure. For each network with more than 20 regions, we concatenated region-wise resting-state fMRI time series across 100 subjects, averaged them, and computed pairwise cosine similarities. We then used agglomerative hierarchical clustering (average linkage) to identify 20 functional subgroups. Within each cluster, time series were averaged to yield a reduced set of representative nodes for modeling; the group-level (consensus) structural connectivity matrices were correspondingly aggregated across the same clusters‡.

As shown in Fig. S2, we recovered a similar pattern of results to those reported in the main text: evidence-weighted prior-variance transformations remained positive and monotonic across all networks with non-zero within-network structural connectivity, and log-Bayes factors comparing structurally informed and uninformed models indicated substantial increases in model evidence, both at the group and subject levels. These results support the generalizability of our approach across atlases.

‡Model inversion was attempted for all 100 subjects using a maximum wall time of 84 hours per job (12 network-specific DCMs per subject). Two jobs did not complete within this limit; as a result, some networks include data from 99 or 98 subjects. These exclusions reflect practical computational constraints rather than model failure or data quality concerns.

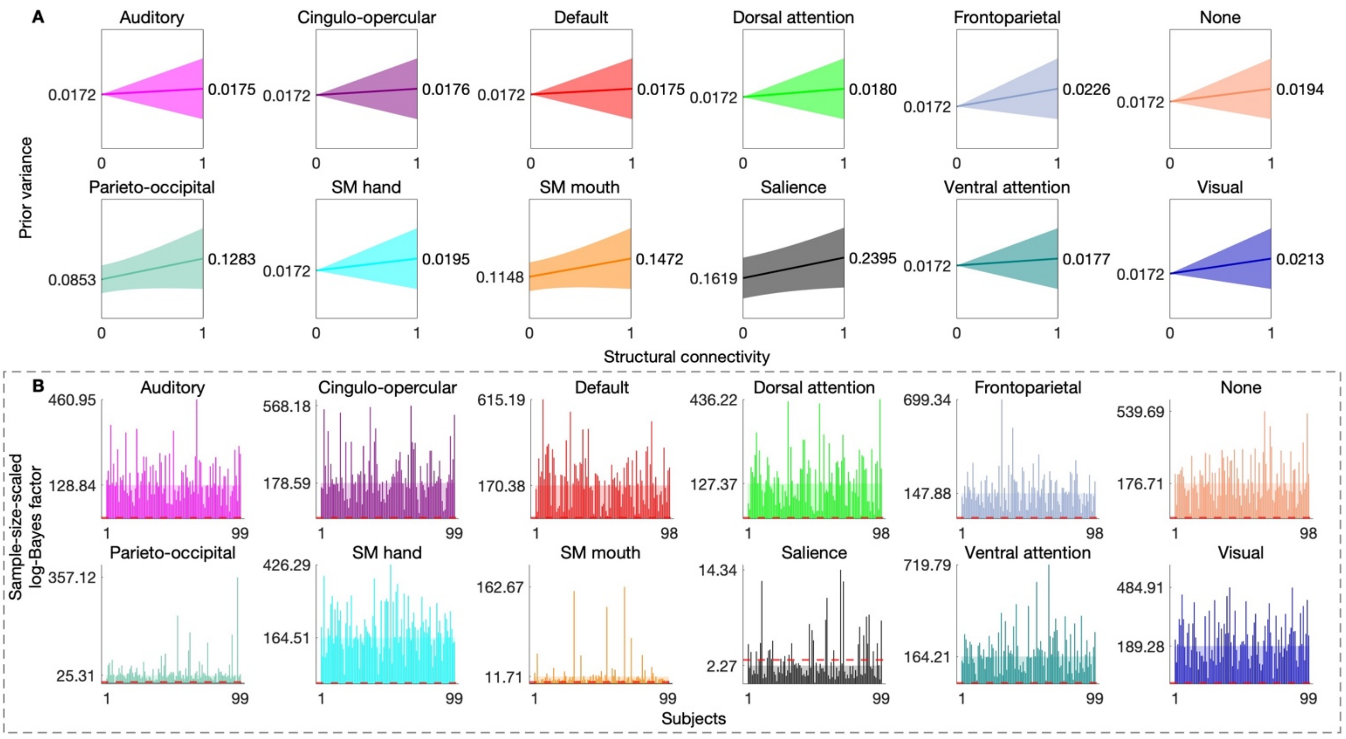

**Fig. S2.** Robustness of structurally informed modeling using the Gordon atlas. To assess the robustness of our findings to parcellation scheme, we repeated the analysis using the Gordon atlas—a functionally derived cortical parcellation. (*A*) Across all 12 networks with non-zero within-network structural connectivity, the evidence-weighted prior-variance transformation remained positive and monotonic. (*B*) Log-Bayes factors comparing hierarchical empirical Bayes models with structure-based priors (derived from evidence-weighted prior-variance transformations) to models with uninformative priors, across 12 effective connectivity brain networks. The semi-transparent bars represent the sample-size-scaled log-Bayes factor for group-level component of the model, while the opaque bars represent the log-Bayes factor for the subject-level component (unaffected by sample-size scaling). The log-Bayes factors indicate substantially greater evidence for models incorporating structure-based priors. Note a dashed red line indicates a log-Bayes factor of 3. Y-axis tick marks indicate the (per-subject) increase in evidence at the group level and the maximum increase in evidence at the subject level, respectively. Fewer than 100 subjects were included in this supplementary analysis due to computational constraints on model inversion. Analyses included SM hand (somatomotor hand area) and SM mouth (somatomotor mouth area) networks.

### Test–retest reliability of structure-based priors

To evaluate the test–retest reliability of our structure-informed hierarchical models, we applied the evidence-weighted prior-variance transformations estimated in the session-1 (test) dataset to the session-2 (retest) dataset. As shown in Fig. S3A, this yielded consistent increases in model evidence across all networks, at both the group and subject levels. Fig. S3B shows that the resulting group-level effective connectivity estimates remained within a plausible range.

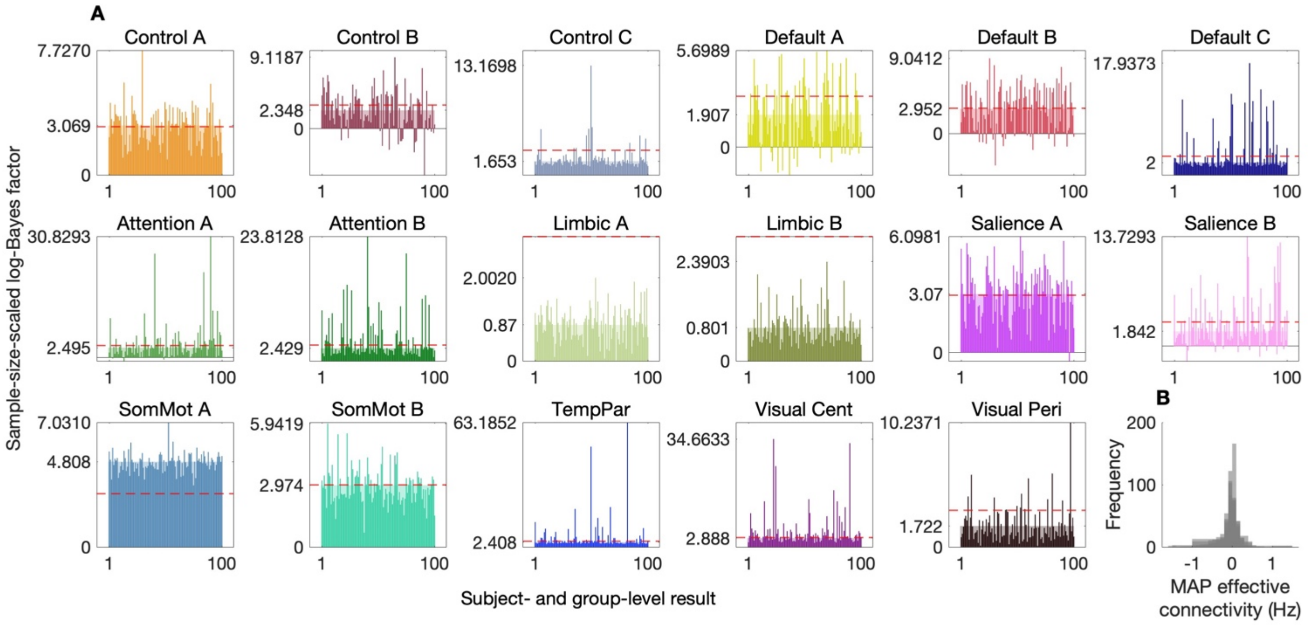

**Fig. S3.** Test-retest reliability of hierarchical empirical Bayes models. (*A*) Log-Bayes factors comparing hierarchical empirical Bayes models with structure-based priors (derived from out-of-session evidence-weighted prior-variance transformations) to models with uninformative priors, across 17 effective connectivity brain networks. The semi-transparent bars represent the sample-size-scaled log-Bayes factor for group-level component of the model, while the opaque bars represent the log-Bayes factor for the subject-level component (unaffected by sample-size scaling). The log-Bayes factors indicate substantially greater evidence for models incorporating structure-based priors. Note a dashed red line indicates log-Bayes factor of 3. Nonzero y-axis tick marks indicate the (per subject) increase in evidence at the group level and the maximum increase in evidence at the subject level, respectively. Networks analyzed include control (Control A–C), default mode (Default A–C), attention (Attention A, B), limbic (Limbic A, B), salience (Salience A, B), somatomotor (SomMot A, B), temporoparietal (TempPar), and visual networks (Visual Cent, central; Visual Peri, peripheral). (*B*) Histogram of maximum *a posteriori* (MAP) group-level effective connectivity estimates for all networks.

### Out-of-sample validation in session-2 dataset

To assess further generalizability, we repeated procedures in which out-of-sample prior-variance transformations were applied to the session-1 fMRI scans for cross-validation subjects (Results, Fig. 4) with the session-2 fMRI scans. As shown in Fig. S4A, structure-informed models again yielded substantial increases in model evidence and resulted in plausible MAP effective connectivity estimates (Fig. S4B).

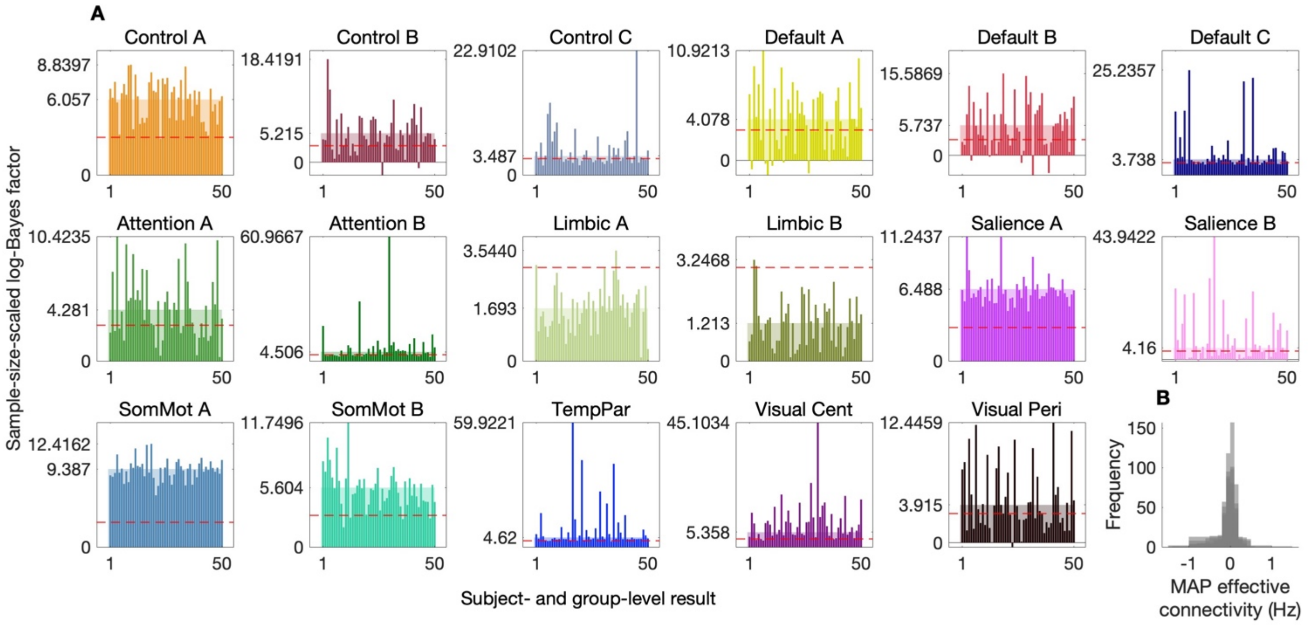

**Fig. S4.** Out-of-sample validation of hierarchical empirical Bayes models (session-2 results). (*A*) Log-Bayes factors comparing hierarchical empirical Bayes models with structure-based priors (derived from out-of-sample evidence-weighted prior-variance transformations) to models with uninformative priors, across 17 effective connectivity brain networks. The semi-transparent bars represent the sample-size-scaled log-Bayes factor for group-level component of the model, while the opaque bars represent the log-Bayes factor for the subject-level component (unaffected by sample-size scaling). The log-Bayes factors indicate substantially greater evidence for models incorporating structure-based priors. Note a dashed red line indicates a log-Bayes factor of 3. Nonzero y-axis tick marks indicate the (per-subject) increase in evidence at the group level and the maximum increase in evidence at the subject level, respectively. Networks analyzed include control (Control A–C), default mode (Default A–C), attention (Attention A, B), limbic (Limbic A, B), salience (Salience A, B), somatomotor (SomMot A, B), temporoparietal (TempPar), and visual networks (Visual Cent, central; Visual Peri, peripheral). (*B*) Histogram of maximum *a posteriori* (MAP) group-level effective connectivity estimates for all networks.

### Permutation-based assessment of structure-based priors

Bayes factors provide a principled means of adjudicating between competing generative models. However, in this hierarchical setting, their magnitude can increase rapidly when model priors become more precise, and those priors happen to more closely resemble the underlying data-generating process (5). This motivates the question: If one were to unknowingly use scrambled structural connectivity in the context of this hierarchical empirical Bayes model, would it yield generalizable evidence gains (like those we have reported)? The permutation analyses reported here suggest not.

To directly evaluate whether the observed improvements in model evidence could be attributed to generic shrinkage (or regularization) effects rather than the specificity of the structure-based prior, we conducted a permutation- and model-comparison-based analysis. Specifically, using BMR (Methods, Eqs. 10–11), we re-evaluated each second-level model used to derive the results shown in Fig. S3 under a randomly permuted (scrambled) version of its prior—preserving equality of the prior variances for $a_{i,j}^{(2)}$ and $a_{j,i}^{(2)}$—while keeping all other model components fixed. This is important because a full *de novo* reinversion under each permuted prior would permit changes in model evidence to reflect not only disruption of the structure-based prior, but also the overall dispersion of second-level RFX (via re-estimation of the second-level precision scaling $\gamma_q^{(2)}$). We then propagated these reduced (scrambled) posteriors to the first level and, again using BMR, computed the corresponding reduced log-evidence for each subject.

For this analysis, we focused on the retest dataset (session 2) DCMs, which constituted the largest non-training set ($S = 100$). For subject $s = 1, \ldots, S$ and model $k$ (with $k = 1$ denoting the structurally informed model, and $k = 2, \ldots, K$ denoting permuted models), we defined the subject-level log-Bayes factor—relative to the corresponding uninformed model—as:

$$\ln \mathrm{BF}^{(1)}_{s,\mathrm{uninformed},k} = F^{(1)}_{s,k} - F^{(1)}_{s,\mathrm{uninformed}}. \qquad \text{[S10]}$$

We assembled $\left\{\ln \mathrm{BF}^{(1)}_{s,\mathrm{uninformed},k}\right\}_{s=1}^{S}$ for each $k$, and performed RFX Bayesian model comparison across models using the standard Dirichlet-multinomial formulation (26, 27), summarizing model prevalence via expected model frequencies and exceedance probabilities $\phi_k$ (the posterior probability that model $k$ is more frequent than any other model in the set). Here, we used a standard routine implemented in the variational Bayesian analysis toolbox (28).

Across networks, the structurally informed model consistently achieved the highest expected model frequency (red bars) and the highest exceedance probability ($\phi_1$), close—with a few exceptions—to one (Fig. S5A). Furthermore, for nearly all networks (as reported in the main text), structurally informed priors yielded positive log-Bayes factors for almost all subjects (Fig. S5B), whereas permuted priors typically did so for fewer than half. One notable exception was the Visual Cent network, where permuted priors more often yielded positive log-Bayes factors; this is consistent with that network being comparatively weakly shaped by the structural-variance mapping (Results, Fig. 5). In sum, scrambled priors did not reproduce the broad, cross-subject evidence gains that underpin our main conclusions, but instead produced gains that were infrequent and idiosyncratic, as expected under chance.

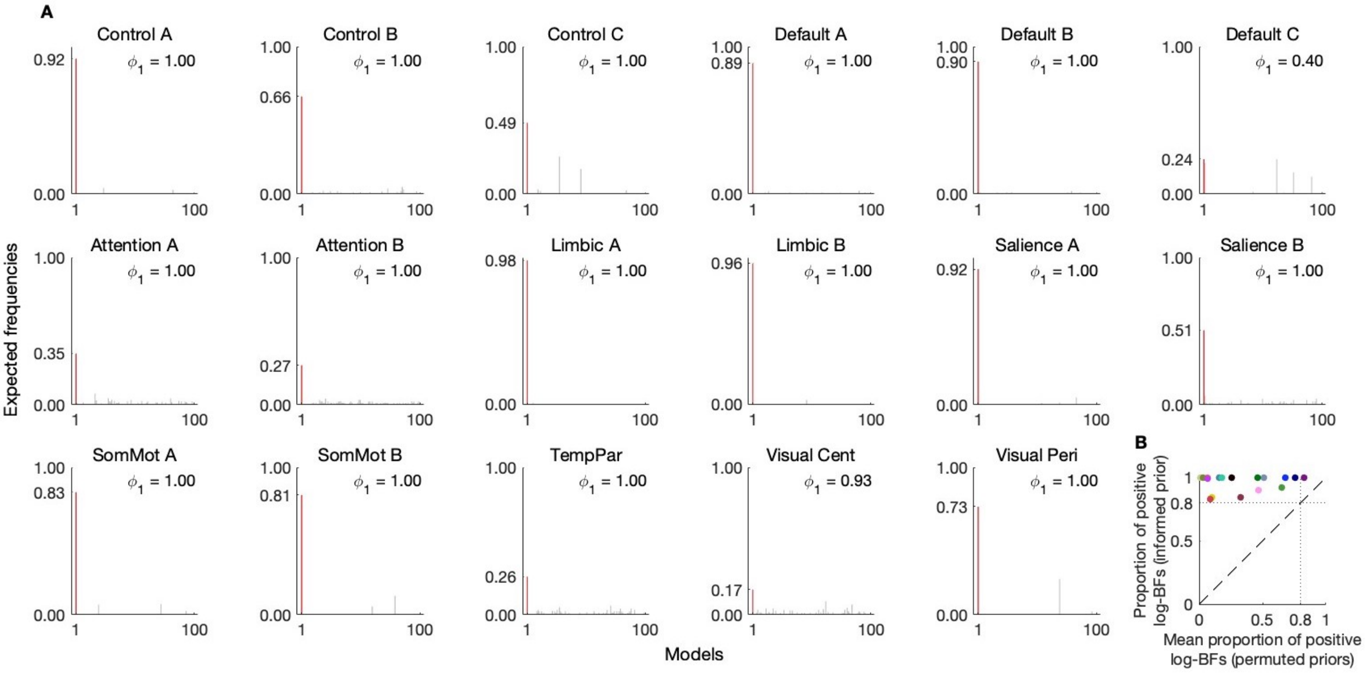

**Fig. S5.** Permutation-based assessment of structure-based priors in 17 brain networks. (*A*) For each network, we performed random-effects Bayesian model comparison over a set of $K = 100$ competing second-level priors: the structurally informed prior ($k = 1$; red bar) and 99 scrambled priors generated by permuting inter-regional prior variances while preserving the overall degree of shrinkage (gray bars). Bars show expected model frequencies and $\phi_1$ denotes the exceedance probability of the structurally informed model (posterior probability that it is more frequent than any alternative). (*B*) Subject-wise prevalence of evidence gains. Points (colored by network) compare the proportion of subjects with positive subject-level log-Bayes factors under the informed prior (y-axis) to the mean proportion under permuted priors (x-axis). The dashed line indicates equality, and dotted lines mark a reference threshold (0.8). Across networks, the structurally informed prior yields broad, cross-subject evidence gains, whereas permuted priors typically yield positive evidence for fewer than half of subjects; Visual Cent shows comparatively higher rates under permutation, consistent with this network being weakly shaped by the structural variance mapping.

### Reliability of effective connectivity estimates

To assess the robustness of our structurally informed hierarchical model, we examined the reliability of group-level effective connectivity estimates. Specifically, we computed pairwise correlations between MAP estimates for each network across datasets. As shown in Fig. S6, effective connectivity estimates were highly consistent across all dataset combinations (all $r \geq 0.66$, $p < 1 \times 10^{-5}$), supporting the stability of the inferred group-level patterns.

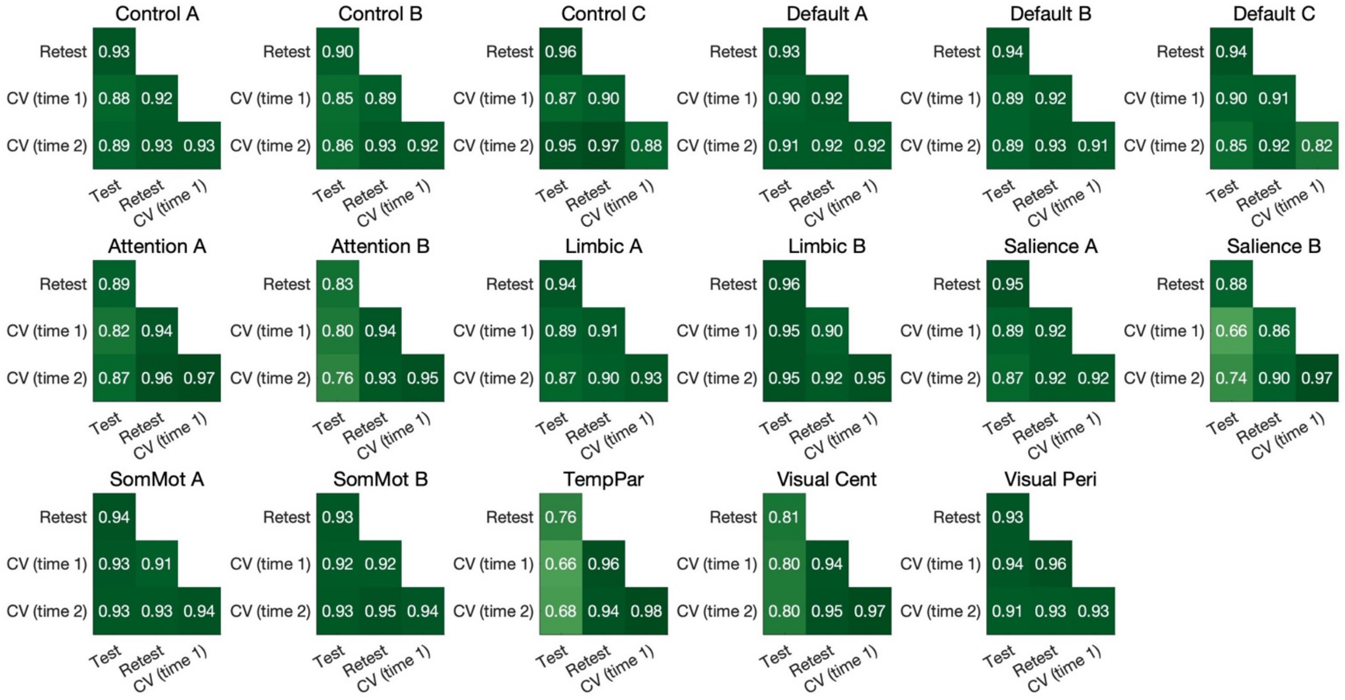

**Fig. S6.** Reliability of effective connectivity estimates across networks and datasets. Pairwise Pearson correlations are shown between group-level maximum *a posteriori* (MAP) effective connectivity estimates derived from the structurally informed hierarchical empirical Bayes model. For each network, estimates were obtained for the four datasets: a test (session 1) and re-test dataset (session 2) of 100 healthy adults, and two cross-validation datasets (CV time 1 and CV time 2) of 50 healthy adults. Correlation values are shown for all pairwise comparisons and reflect high reliability (all $r \geq 0.66$, $p < 1 \times 10^{-5}$).
